# Accommodating site variation in neuroimaging data using normative and hierarchical Bayesian models

**DOI:** 10.1101/2021.02.09.430363

**Authors:** Johanna M. M. Bayer, Richard Dinga, Seyed Mostafa Kia, Akhil R. Kottaram, Thomas Wolfers, Jinglei Lv, Andrew Zalesky, Lianne Schmaal, Andre Marquand

## Abstract

The potential of normative modeling to make individualized predictions from neuroimaging data has enabled inferences that go beyond the case-control approach. However, site effects are often confounded with variables of interest in a complex manner and can bias estimates of normative models, which has impeded the application of normative models to large multi-site neuroimaging data sets. In this study, we suggest accommodating for these site effects by including them as random effects in a hierarchical Bayesian model. We compared the performance of a linear and a non-linear hierarchical Bayesian model in modeling the effect of age on cortical thickness. We used data of 570 healthy individuals from the ABIDE (autism brain imaging data exchange) data set in our experiments. In addition, we used data from individuals with autism to test whether our models are able to retain clinically useful information while removing site effects. We compared the proposed single stage hierarchical Bayesian method to several harmonization techniques commonly used to deal with additive and multiplicative site effects using a two stage regression, including regressing out site and harmonizing for site with ComBat, both with and without explicitly preserving variance related to age and sex as biological variation of interest. In addition, we made predictions from raw data, in which site has not been accommodated for. The proposed hierarchical Bayesian method showed the best predictive performance according to multiple metrics. Beyond that, the resulting z-scores showed little to no residual site effects, yet still retained clinically useful information. In contrast, performance was particularly poor for the regression model and the ComBat model in which age and sex were not explicitly modeled. In all two stage harmonization models, predictions were poorly scaled, suffering from a loss of more than 90 % of the original variance. Our results show the value of hierarchical Bayesian regression methods for accommodating site variation in neuroimaging data, which provides an alternative to harmonization techniques. While the approach we propose may have broad utility, our approach is particularly well suited to normative modelling where the primary interest is in accurate modelling of inter-subject variation and statistical quantification of deviations from a reference model.

**Highlights:**

- Development and presentation of normative modeling approach based on hierarchical Bayesian modeling that can be applied to large multi-site neuroimaging data sets.
- Comparison of performance of Hierarchical Bayesian model including site as predictor to several common ways to harmonize for multi-site effects.
- Presentation of normative modeling as site correction tool.

## 2 Introduction

The most prominent paradigm in clinical neuroimaging research has for a long time been case-control approaches which compare averages of groups of individuals on brain imaging measures. Case-control inferences can be clinically meaningful under some circumstances when the group mean is a good representation of each individual in the group. However, this pre-condition has been challenged recently, demonstrating that the biological heterogeneity within clinical groups can be substantial (Marquand et al., 2016). For example, the structure and morphology of the brain have been found to vary between individuals in dynamic phases like adolescence (Foulkes and Blakemore, 2018) and within clinical groups, such as bipolar disorder and schizophrenia (Wolfers et al., 2018a) and attention deficit disorder (Wolfers et al., 2019). In addition, inter-individual differences have shown to not necessarily be in line with results obtained via the group comparison approach (Wolfers et al., 2019). Such heterogeneity has been considered a potential cause for the lack of differences between clinical groups and controls within the standard group comparison approach (Feczko et al., 2019) and the failure to replicate findings between studies (Fried, 2017). As a consequence, there has been a shift in focus towards taking into account variation at the individual level (Marquand et al., 2019). This is in line with a trend towards personalized medicine or “precision medicine” (Mirnezami et al., 2012), where characteristics of an individual are used to guide the treatment of mental disorders.

This shift has been accompanied by a trend towards approaches that go beyond comparing averages of distinctly labeled groups (Insel et al., 2010; Insel, 2014). Among them, normative modeling has been successfully used to capture inter-individual variability and make predictions at the individual level. The strength of normative modeling lies within the ability to map variation across one or more biological response variables (e.g., brain volume) onto one or more covariates or predictor variables (e.g., age), redefining the variation in the first dimension as explained by this new covariate(s) of interest. This concept allows to describe the normative variation, thus the range containing e.g., 95 % of all individuals, as a function of the covariates. As a consequence, each individual’s score in considered relation to the variation in the reference group defined by the covariates, allowing to calculate a z-score of deviation from the norm at the level of the individual. The concept is similar to the use of growth charts in pediatric medicine, in which height and weight are expressed as a function of age. Hence, in this setting, an individual’s height or weight is not considered by its absolute value, but expressed as a percentile score of deviation fluctuating with age, with the median line corresponding to the 50% percentile and defining the norm, or average height.

In neuroimaging, normative models have been applied to clinical and non-clinical problems using various covariates, statistical modeling approaches (for an overview see (Marquand et al., 2019, 2016)) and targeting a variety of response variables. In general, any variable can be used as a covariate in a normative model targeting neuroimaging measures, as long as the variation along the co-varying dimension is not zero. However, normative models with age and sex as covariates and brain volume as response variable are currently more frequently found in the literature (Wolfers et al., 2020, 2018b; Zabihi et al., 2019a; Kessler et al., 2016). These implement the growth charting idea applied to high dimensional brain imaging data. For example, a normative model of a brain structure can be created based on the variation of individuals in population based cohorts. The estimated norm can be used to infer where individuals with clinical symptoms can be placed with respect to the reference defined by the normative model. This has been the recipe of many recently published studies using the normative modeling framework (Wolfers et al., 2020; Bethlehem et al., 2018; Wolfers et al., 2019; Lv et al., 2020). Underlying this approach is the assumption that the individually derived patterns of deviation uncover associations to clinical/behavioral variables that would be obscured by averaging across groups of individuals. However, the amount of data necessary to create normative models poses a challenge to normative modeling in neuroimaging, as the cost and time factor associated with neuroimaging data impedes the collection of large neuroimaging samples in a harmonized way. One exceptional example, where large scale data collection succeeded and included both harmonized scanners and scanning protocols, is the UK Biobank initiative. When launched in 2006, it aimed to scan 100,000 individuals at four different scanning locations [https://www.ukbiobank.ac.uk/explore-your-participation/contribute-further/imaging-study](Miller et al., 2016). Other neuroimaging initiatives have also taken on the challenge to collect neuroimaging data in large scale quantities and have relied on harmonized scanning protocols, but did not collect the data using harmonized scanners (i.e., ADNI (Mueller et al., 2005) and the ABCD study (Volkow et al., 2018)). Nonetheless, the restricted age ranges (e.g., 40-69 years in UK Biobank (Miller et al., 2016)), or focus on a particular (clinical) cohort (e.g., Alzheimer’s in ADNI, (Mueller et al., 2005)) limit their utility for estimating normative models mapping the normative association between, for example, age and brain structure or function.

An alternative way to obtain large neuroimaging datasets and assess data from a large number of subjects is by pooling or sharing data that has already been collected. One example is the Enhancing NeuroImaging and Genetics through Meta-Analysis (ENIGMA) consortium (Thompson et al., 2020). ENIGMA succeeded in pooling neuroimaging and genetics data of thousands of individuals, including healthy individuals and individuals with psychiatric or neurological disorders. The strategy of data sharing initiatives like ENIGMA is to collect already collected data from different cohorts and different scanning sites and harmonize preprocessing and statistical analysis with standardized protocols. However, a major disadvantage is the presence of confounding “scanner effects” (Fortin et al., 2017, 2018) (e.g., differences in field strength, scanner manufacturer etc. (Han et al., 2006))). These confounding effects present as site correlated biases that cannot be explained by biological heterogeneity between samples. An example of those effects on derived measures of cortical thickness can be found in Fig. 1a. They result from a potentially complex interaction between site and variables of interest, manifesting in biases on lower and higher order properties of the distribution of interest, such as differences in mean and standard deviations, skewness and spatial biases Fig. (1a, 1b), and cannot be explained by e.g., differences in age or sex Fig. (1c). As the origin of these effects might not only be related to the *scanner per se*, but extend to various factors related to a single acquisition site (Gronenschild et al., 2012), we will refer to them as *site effects* from here on.

**Figure 1:**
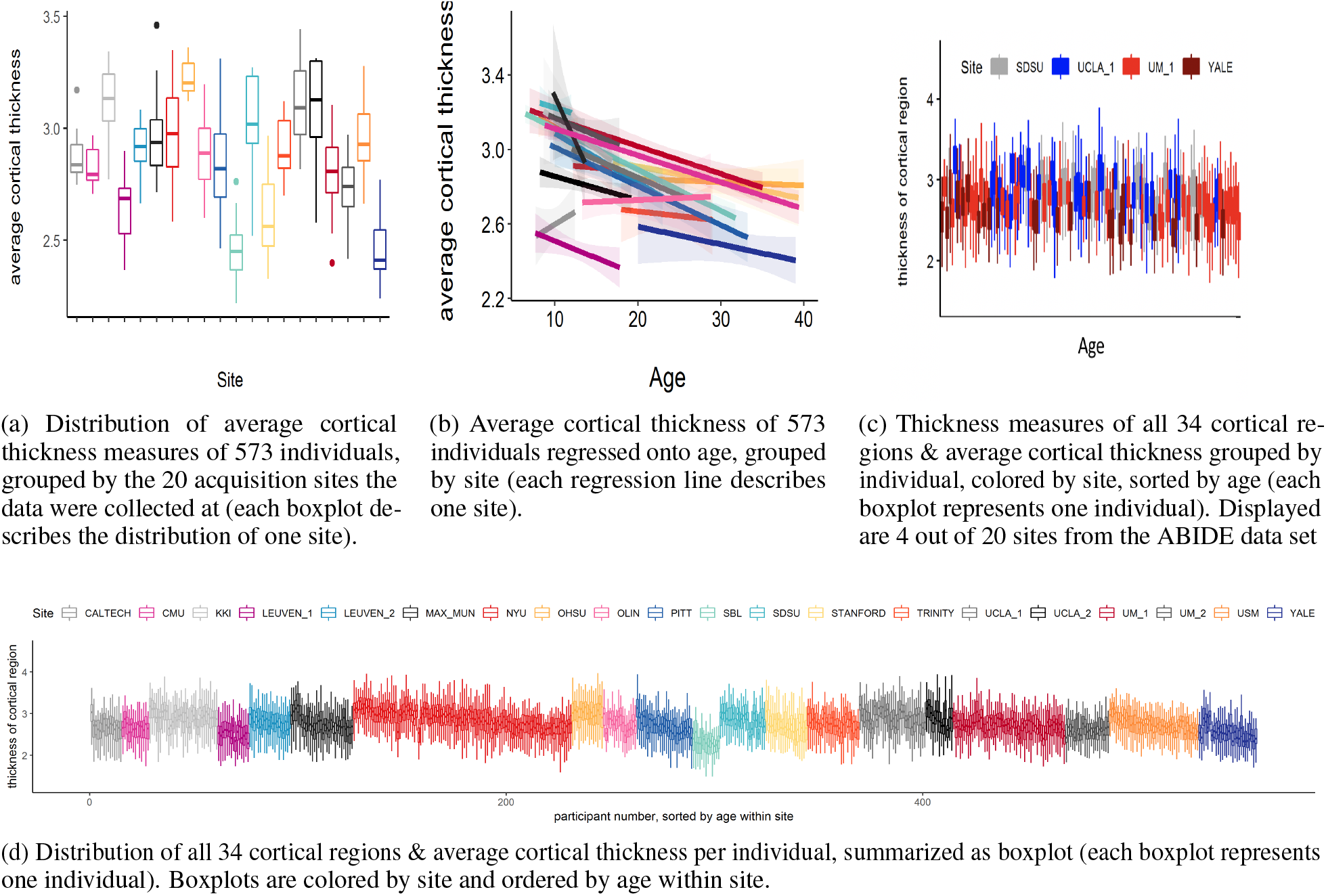
Site effects in 573 healthy individuals from the ABIDE data set.

As outlined in the previous paragraph, the effort to create large samples to capture between subject variability often induces site-driven variability. This issue of site-driven variability in shared neuroimaging data has been acknowledged and has led to the development of harmonization methods at a statistical level. A common approach to deal with site effects is through “harmonizing” by, e.g., confound regression. One example of this approach is a set of algorithms summarized under the name “ComBat” (Fortin et al., 2017). The method had originally been developed by Johnson et al. (2007), who used empirical Bayes to estimate “batch effects”, referring to non-biological variation in micro-array experiments for gene expression data. Fortin and colleagues adapted the framework to apply to neuroimaging data (Fortin et al., 2017). In ComBat, additive and multiplicative site effects on a particular target unit (e.g., a particular brain voxel for one participant) are estimated using empirical Bayes and by placing a prior distribution over estimates for these units. The estimate of the scanner effect is then used to adjust the prediction. Newer versions also allow to preserve variance of interest in the model, for example for age, sex or diagnosis (Fortin et al., 2017, 2018). ComBat has been applied to several types of neuroimaging data, including diffusion tensor imaging data (DTI, (Fortin et al., 2017)) and structural magnetic resonance imaging data, in particular measures of cortical thickness (Fortin et al., 2018).

The estimation and interpretation of data harmonization strategies is relatively straightforward if site effects are orthogonal to the effect of interest and uncorrelated with other covariates in the model (Chen et al., 2014). However, this is often not the case in neuroimaging cohorts containing data pooled from several sites. For example, many individual neuroimaging samples are restricted to a specific age range, leading to age being correlated with site effects. In this scenario, removing an estimate of the scanner effect can also remove (biological) variation that would be of interest. This is undesirable from the point of view of normative modelling, where analysis focuses on deviations from a reference model because the deviations from a model where scanner effects have been regressed out in a two stage procedure may be different from a model that integrates site information in a single regression. Data harmonisation also has other features that make it suboptimal for normative modelling: first, many implementations pool estimates over voxels or parcels, which means that the centiles of population variation at a particular brain region are influenced by other regions. Second, a problem that is not widely appreciated in the neuroimaging field is that data harmonisation strategies can yield overly optimistic inferences in downstream analyses due to an under-estimation of the uncertainty of model coefficients Nygaard et al. (2016)

For these reasons, we suggest an alternative approach to deal with site effects in neuroimaging data. While we focus in particular on normative modeling, our approach can be also applied to other neuroimaging data analyses scenarios. We propose a hierarchical Bayesian approach in which we include site as a random effect in the model, avoiding the exclusion of meaningful variance correlated with site by predicting site effects as part of the model instead of removing them from the data.

This approach is similar to the approach by Kia et al. (2020), who used hierarchical Bayesian regression (HBR) in a similar way for multi-site normative modeling in a pooled neuroimaging data set, which contained 7499 participants who were scanned with 33 different scanners. Kia et al. (2020)’s estimate of site variation is based on a partial pooling approach, in which the variation between site-specific parameters is bound by a shared prior. The approach showed better performance when evaluated with respect to metrics accounting for the quality of the predictive mean and variance compared to a complete pooling of site parameters and to ComBat harmonization, and similar performance to a no-pooling approach, with the benefit of reduced risk of over-fitting due to the shared site variance. Moreover, Kia et al. (2020) also showed that the posterior distribution of site parameters from the training set can also be used as an informed prior to make predictions in an unseen, new test set, outperforming predictions from complete pooling and uninformed priors. The method was also able to preserve and parse heterogeneity between individuals with varying clinical diagnoses in associated brain regions of 1017 clinical patients of the study.

The present paper is a replication and extension of the approach by Kia et al. (2020). Based on several successful attempts of using Gaussian Process Regression to map non-linearity in normative models (Kia and Marquand, 2018; Marquand et al., 2016, 2014), we extend the normative model with the capacity to account for site effects by adding a Gaussian process to model non-linear effects between age and the brain structure. In addition, our model is fully Bayesian and entails a hierarchical structure, including priors and hyper-priors for each parameter. We used the healthy control sample from the ABIDE (autism brain imaging data exchange, http://preprocessed-connectomes-project.org/abide/) (Di Martino et al., 2014) data set to compare a non-linear, Gaussian process version of the model, to a linear hierarchical Bayesian version accounting for site effects that does not include the Gaussian Process term. We show that the hierarchical Bayesian models including a site parameter perform better than existing methods for dealing with additive and multiplicative site effects, including *ComBat* and regressing out site. Subsequently, we validate the hierarchical Bayesian models in the autism sample of the ABIDE data set and test their ability to retain clinically useful variance while correcting for site effects. We discuss the normative hierarchical Bayesian methods with regard to their implications for neuroimaging data-sharing initiatives and their use as general technique to correct for site effects.

## 3 Methods

In this section, we will introduce the data used in this study and the pre-processing steps applied, followed by a conceptual and mathematical description of our approach to include site as predictor in a normative hierarchical Bayesian model. We will also illustrate other methods (than including site as predictor) to accommodate for site effects that will be used to validate our approach against. Lastly, we will outline which measures will be used for model comparison.

### 3.1 Data

The following sub-section aims to give a description of the ABIDE data set, including a study on the scope of site effects in the data.

#### 3.1.1 ABIDE data set

The ABIDE consortium (http://preprocessed-connectomes-project.org/abide/) was founded to facilitate research and collaboration on autism spectrum disorders by data aggregation and sharing. The consortium provides a publicly available structural magnetic resonance imaging (MRI) data set and corresponding phenotypic information of 539 individuals with autism spectrum disorder and 573 age-matched typical controls. For this study, we used 569 controls for development and performance testing of the models, out of which 470 were male. In a subsequent step, we applied the hierarchical Bayesian models to 482 individuals with autism from the same data set, out of which 430 were male. The data were processed using a standardized protocol (Craddock et al., 2013) of the FreeSurfer standard pipeline (Desikan-Kiliany Atlas) as part of the *Preprocessed Connectomes Project* (Craddock et al., 2013) and has been made available for download on the *preprocessed* section of the ABIDE initiative. For the current study, we focused on cortical thickness measures of the 34 bilateral regions (averaged between left and right hemisphere) of the Desikian Killiany atlas parcellation (Desikan et al., 2006) as a part of the FreeSurfer (Fischl et al., 2004) output and the average cortical thickness across all 34 regions. We chose to include only cortical thickness measures since they show a strong (negative) association with age (unlike measures of surface area, which remain more stable across the lifespan (Storsve et al., 2014)).

#### 3.1.2 Site effects in the ABIDE data set

The ABIDE data set has been obtained by aggregating data from 20 independent samples collected at 17 different scanning locations (Di Martino et al., 2014). Although all data has been collected with 3 Tesla scanners and preprocessed in a harmonized way (Craddock et al., 2013), sequence parameters for anatomical and functional data, as well as type of scanner varied across sites (Di Martino et al., 2014). In addition, sites differ in distribution of age and sex and in sample size. An overview of site-specific data is provided in Table 1 and in (Di Martino et al., 2014).

**Table 1:** The scanner parameters and sample specifications of the ABIDE data set.

| Site<br>(Abbreviation) | Manufacturer | Platform | Voxel Size<br>[mm] | TR<br>[ms] | TE<br>[ms] | n | males<br>[%] | age range<br>[years] |
| --- | --- | --- | --- | --- | --- | --- | --- | --- |
| CALTECH<sup>a</sup> | SIEMENS | TIM TRIO | 1.0×1.0×1.0 | 1590 | 2.73 | 15 | 0.73 | 17-39 |
| CMU<sup>b</sup> | SIEMENS | TIM TRIO | 1.0×1.0×1.0 | 1870 | 2.48 | 13 | 0.77 | 20-40 |
| KKI<sup>c</sup> | Philips | Achieva | 1.0×1.0×1.0 | 8 | 3.7 | 33 | 0.73 | 8-12 |
| LMU<sup>d</sup> | SIEMENS | VERIO | 1.0×1.0×1.0 | 1800 | 3.06 | 15 | 1 | 18-29 |
| NYU<sup>e</sup> | SIEMENS | ALLERGRA | 1.3×1.0×1.3 | 2530 | 3.25 | 20 | 0.75 | 12-16 |
| OLIN<sup>f</sup> | SIEMENS | ALLEGRA | 1.0×1.0×1.0 | 2500 | 2.74 | 30 | 0.9 | 7-35 |
| OHSU<sup>g</sup> | SIEMENS | TIM TRIO | 1.0×1.0×1.0 | 2300 | 3.58 | 105 | 0.75 | 6-31 |
| SDSU<sup>h</sup> | GE | MR750 | 1.0×1.0×1.0 | 11.08 | 4.3 | 15 | 1 | 8-12 |
| SBL<sup>i</sup> | Philips | INTERA | 1.0×1.0×1.0 | 9 | 3.5 | 16 | 0.875 | 10-23 |
| STANFORD<sup>j</sup> | GE | SIGNA | 0.859 × 1.500 × 0.859 | 8.4 | 1.8 | 27 | 0.85 | 9-33 |
| TRINITY<sup>k</sup> | Philips | Achieva | 1.0 × 1.0 × 1.0 | 3.9 | 8.5 | 13 | 1 | 20-39 |
| UCLA 1<sup>l</sup> | SIEMENS | TIM TRIO | 1.0 × 1.0 × 1.2 | 2300 | 2.84 | 22 | 0.73 | 8-16 |
| UCLA 2<sup>m</sup> | SIEMENS | TIM TRIO | 1.0 × 1.0 × 1.2 | 2300 | 2.84 | 20 | 0.8 | 7-12 |
| LEUVEN 1<sup>n</sup> | Philips | Achieva | 0.98 × 0.98 × 1.20 | 9.6 | 4.6 | 25 | 1 | 12-25 |
| LEUVEN 2<sup>o</sup> | Philips | Achieva | 0.98 × 0.98 × 1.20 | 9.6 | 4.6 | 32 | 0.88 | 9-17 |
| UM 1<sup>p</sup> | GE | SIGNA | 1.2 × 1 × 1 | 250 | 5.7 | 13 | 0.85 | 9-13 |
| UM 2<sup>q</sup> | GE | SIGNA | 1 × 1 × 1.2 | 250 | 5.7 | 54 | 0.69 | 8-19 |
| PITT<sup>r</sup> | SIEMENS | ALLERGRA | 1.1×1.1×1.1 | 2100 | 3.93 | 21 | 0.95 | 13-28 |
| USM<sup>s</sup> | SIEMENS | TIM TRIO | 1.0×1.0×1.2 | 2300 | 2.91 | 43 | 1 | 8-39 |
| YALE<sup>t</sup> | SIEMENS | TIM TRIO | 1.0×1.0×1.2 | 1230 | 1.73 | 28 | 0.71 | 7-17 |

<sup>a</sup>Caltech Institute of Technology; <sup>b</sup>Carnegie Mellon University; <sup>c</sup>Kennedy Krieger Institute; <sup>d</sup>Ludwig Maximilians University Munich; <sup>e</sup>NYU Langone Medical Center; <sup>f</sup>Olin Institute of Living at Hartford Hospital; <sup>g</sup>Oregon Health and Science University; <sup>h</sup>San Diego State University; <sup>i</sup>Social Brain Lab; <sup>j</sup>Stanford University; <sup>k</sup>Trinity Centre for Health Sciences; <sup>l</sup>University of California Los Angeles 1; <sup>m</sup>University of California Los Angeles 2; <sup>n</sup>University of Leuven 1; <sup>o</sup>University of Leuven 2; <sup>p</sup>University of Michigan 1; <sup>q</sup>University of Michigan 2; <sup>r</sup>University of Pittsburgh School of Medicine; <sup>s</sup>University of Utah School of Medicine; <sup>t</sup>Yale Child Study Center

The ABIDE data set is affected by site specific effects that are unlikely to be explained by biological variation. They manifest as linear and non-linear interactions between *scanning site*, covariates (for example age and sex), and cortical measures. Similar to *batch effects* in genomics (Leek et al., 2010), those effects lead to a clustering of the data caused by external factors related to the scanning and analysis process. With the aim to estimate to which extent the ABIDE data set is affected by site effects, we calculated an ANCOVA with age as covariate in the healthy control sample of the data set. It revealed that average cortical thickness differed between site (main effect site: F(19, 516) = 4.4, p< 0.1 ×10^−8^, sum contrast). In addition we tested for differences in variance between sites. Bartlett’s sphericity test Bartlett (1937) showed a difference in variance between sites even after regressing out variance that could be explained by age and sex (p < 0.001). The site effects in the healthy control sample of the ABIDE data set are visualized in Fig. 1.

### 3.2 Pre-processing of the ABIDE data set

Measures of cortical thickness were extracted from the arpac.stats files as part of the Freesurfer output of 1051 individuals in the ABIDE data set, separately for left and right hemisphere. As a first step, the measures were scanned for outliers. The criterion applied to mark a value as an outlier was if it was above or beyond 2 inter quartile ranges from the mean of all values for that region and hemisphere. This quite liberal criterion was applied with the aim to detect not outliers in a mathematical sense (+/-95 % confidence interval), but to detect impossible values. This leads to the removal of 1055 out of 162905 data points (0.006%) of all values. After this step, the values of right and left hemisphere for each region were averaged (in case the value of one hemisphere was missing, the value of the remaining hemisphere was considered to be the average.) This procedure was preformed including *all* participants (control sample and autism sample, per region).

### 3.3 Splitting the ABIDE data set into training and test sets

To evaluate the performance of the models, we split the the healthy control data set into a training set (70% of data, n=389) and a test set (30% of data, n=166) using the R package *caret* and *splitstackshape*, while the distribution of age, sex and site was preserved between sets. Thus, training and test sets contained individuals from the same sites (“within-site-split”). For the clinical autism set, information from all individuals with autism that survived outlier correction (n=482) were used. Subsequently, the control training and both the control and clinical autism test sets were standardized region-wise based on location and scale parameters of the training set. For the model estimation process only complete pairs of observations (per region) were used. An overview of the distribution of age and sex for the training and test sets for healthy controls and individuals with autism can be found in Fig. 2

**Figure 2:**
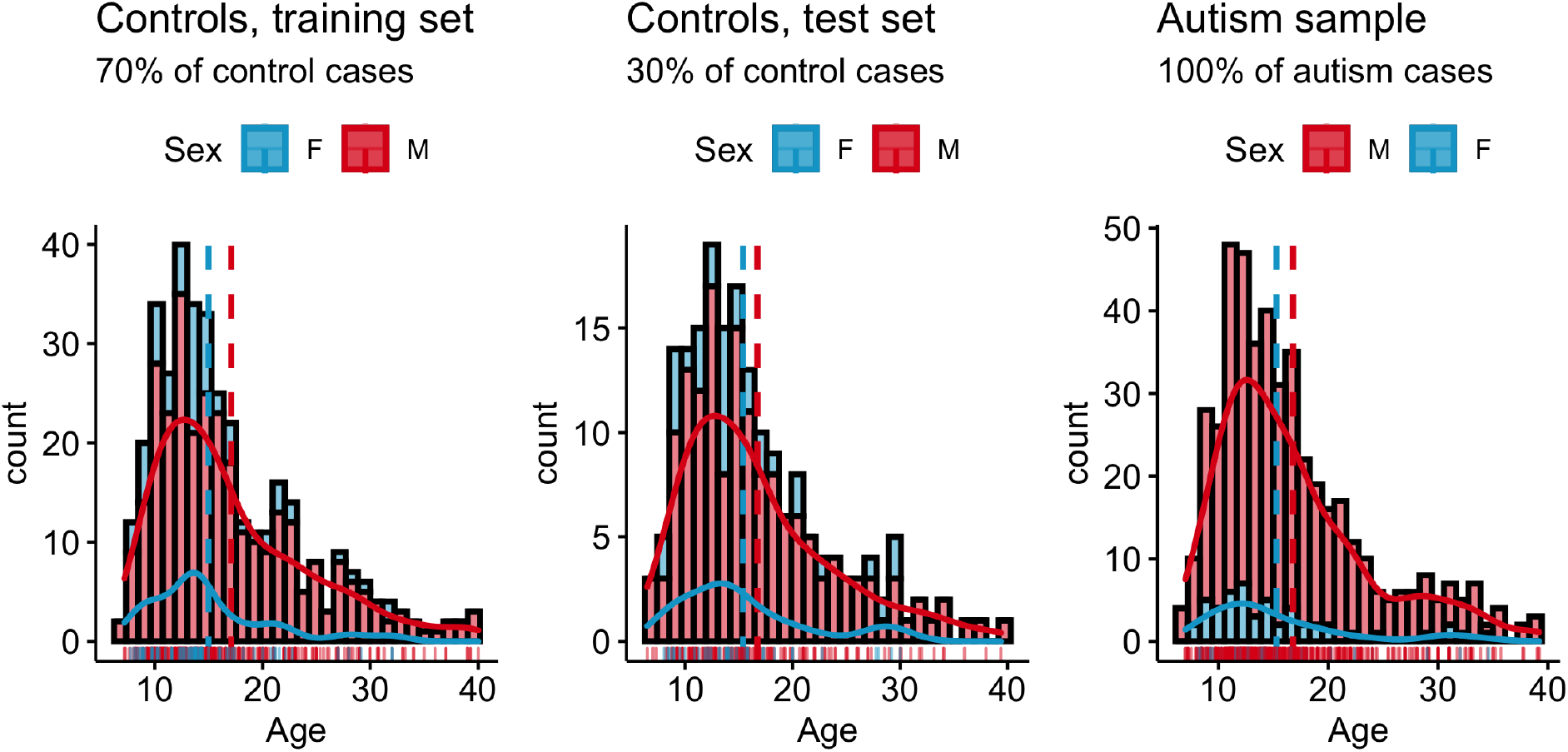
Overview over phenotypic information in the ABIDE data set. Controls: Age male subjects: *M* = 17.5., *SD* = 8.3. Age female subjects: *M* = 15.6, *SD* = 7.0., range = 6.5-40; **Autism sample**: Age male subjects: *M* = 16.9., *SD* = 6.5. Age female subjects: *M* = 15.1, *SD* = 5.8., range = 8-39;

### 3.4 Site as a predictor in a Hierarchical Bayesian Model

With the aim to create reliable normative models in multi-site neuroimaging data, we developed and compared two versions of a hierarchical Bayesian models that include site as a predictor. In a hierarchical linear version of the model, site is modeled hierarchically, resulting in a random effect for site (“Hierarchical Bayesian Linear Model, HBLM”). In a non-linear version of the model, a Gaussian Process for age is added to test whether performance is increased if the model is also able to capture non-linear effects between age and *thickness of the cortical region* (“Hierarchical Bayesian Gaussian Process Model, HBGPM”). Both hierarchical Bayesian models were trained and tested in a within-site split (see section 3.3 on splitting the multi-site ABIDE data set.)

### 3.5 Comparison models

To get a better understanding of the performance of our approach, we performed a second analysis, comparing the hierarchical Bayesian approach with site as predictor to predictions made from a data that other methods managing site effects had been applied on. In the following, those alternative models will be summarized under the term *comparison models*.

Of note, the approach used to accommodate for site effects in the comparison models is fundamentally different from the approach used in the hierarchical Bayesian models. In the hierarchical Bayesian approach, multi-level modeling is used to account for site-variance without removing it, whereas different methods of *harmonization* are used on the data to remove variance related to site as part of the comparison models approach.

In detail, the comparison model approach entailed a two-step procedure, in which site effects are first *harmonized* by three different common models of site harmonization, and then a simple Bayesian linear algorithm, with an additive term for age and sex, but without site as a predictor is used to make predictions in Stan (Stan Development Team, 2020b). The harmonization procedures include i) regressing out site effect from the cortical thickness measures using linear regression and using the residuals as input to the simple Bayesian linear model (thus, removing additive variant components of site), ii) using ComBat (Johnson et al., 2007; Fortin et al., 2017, 2018) to clear the data from site effects (thus, harmonizing for additive and multiplicative effects of site), and iii) using ComBat as above, but explicitly preserving the variance associated with sex and age; an approach which will be referred to as *modified ComBat* in the following. Predictions made from raw data (thus, without any treatment of site effects) were used as a baseline model (Fig. 3).

**Figure 3:**
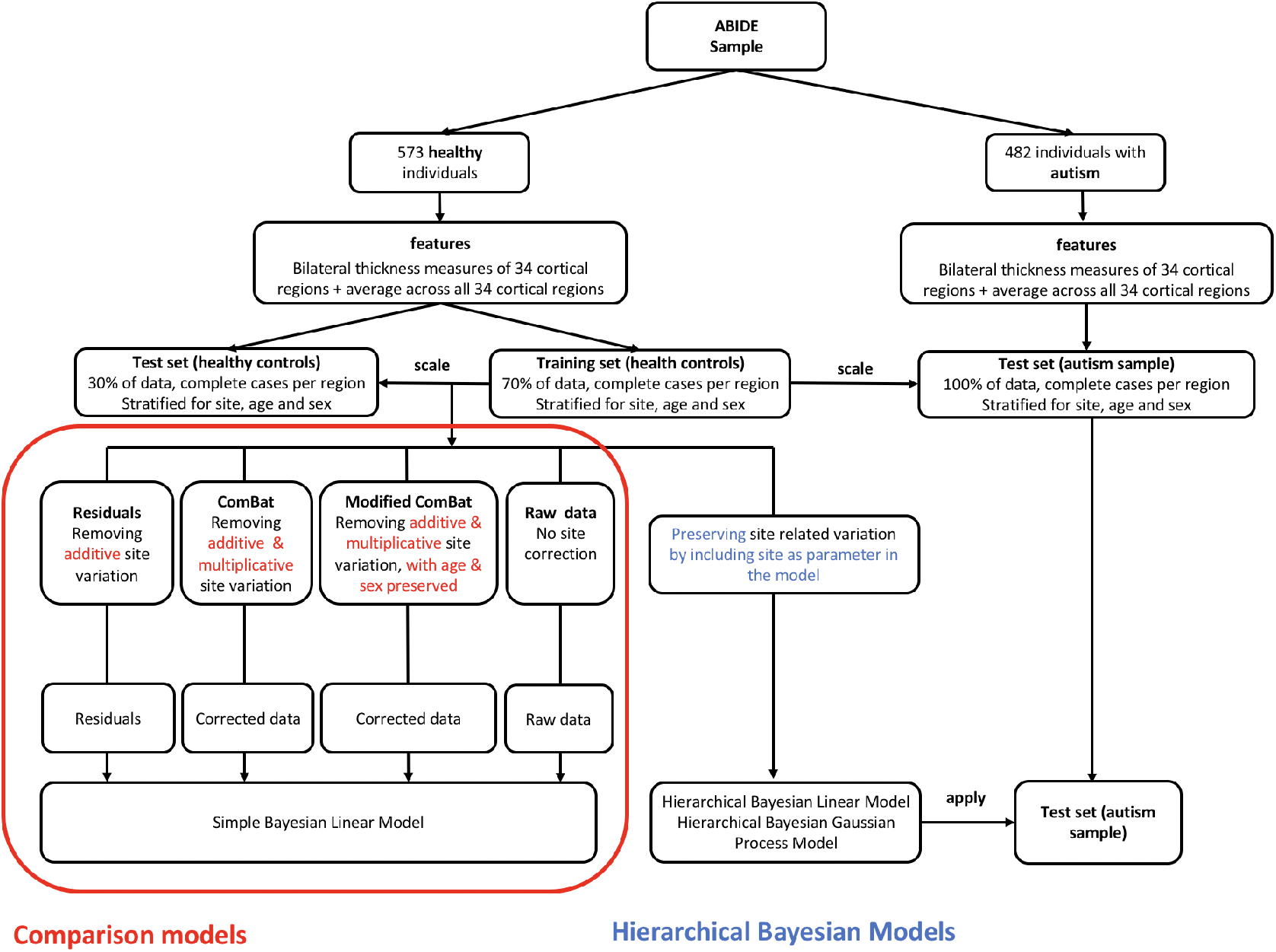
Pipelines for hierarchical Bayesian and comparison models.

### 3.6 Impact of models on variation of interest

Beyond that, we conducted a third step in which we tested the ability of our models to be used as a site correction tool. For any model to be used for site correction, it is not only desirable to correct for site, but at the same time to retain non-site related variation of interest. Thus, in this step we applied the hierarchical Bayesian models that had been trained on the training set of healthy controls to the autism sample of the ABIDE data sets, comparing their results to those of the test controls. This direct comparison between out of sample controls and out of sample clinical data allows for an accurate estimate of distortions of the model on variation of interest. An overview over all pipelines for all models can be found in Fig. 3.

### 3.7 Performance measures

#### 3.7.1 Measures of model performance

Model performance is assessed using several common performance metrics. The Pearson’s correlation coefficient *ρ* indicates the linear association between true and predicted value of cortical thickness measures. However, correlations are not a sensitive error measure and cannot capture the mismatch between true and predicted value. Hence, we also calculate the standardized version of the root mean squared error (SRMSE) and the point-wise log-likelihood at each data point in the test set as a metric indicating deviance from the true value. However, these measures only take into account the estimate of the mean, and do not account for variations in the estimate of the variance. Thus, we also compute the proportion of variance explained (EV) by the predicted values and a standardized version of the log-loss (mean standardized log-loss, MSLL (Rasmussen and Williams, 2006)). The latter does not only take into account the variance of the test set, but also standardizes it by the variance of the training set, making a comparison between the models possible. This step is necessary as various methods of correcting for site might also have an impact on the variance remaining in the data.

#### 3.7.2 Measures of goodness of the simulation in Stan

Parameters indicating the goodness of the model simulation process in Stan itself, like convergence, effective sample size, and trace plots can be found in the supplementary material.

### 3.8 Model specification

In this section, we show how normative models describing the association between age, sex, and cortical thickness measures can be modeled on data comprising site effects using a hierarchical Bayesian linear mixed model with a Gaussian Process term, which allows to model non-linear association between age and cortical thickness measures. Following the notations of (Gelman, 2008; Rasmussen and Williams, 2006), we model a target vector **y** ∈ ℝ^*n*×1^ containing the the individual responses *y*_*i*_ for each subject *i* = 1,. .., *n* and each region, using a latent function **f** = *f*(**x**). *f*_*i*_ = *f*(x_*i*_) is the evaluation of the latent function for an input vector **x**_*i*_ containing all *p* input variables of subject *i*, and is considered to differ from the true response variables by additive noise *ϵ*_*i*_ with the variance *η*_*i*_ and *N*(0, σ^2^) along the diagonal, with **I** being an *n* × *n* identity matrix:

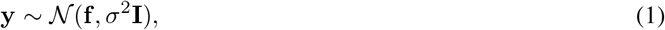

or, for the individual case: with:

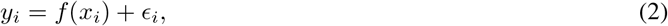

with:

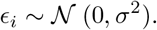

The ability of the model to deal with site effects is obtained by introducing a random effect for site *s* = 1, 2,. .., *q* so that the prediction for the *i*^*th*^ subject is a combination of fixed and varying effects:

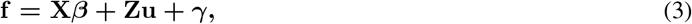

where *γ* is an additional non-linear component (defined in (5) below) and the estimate for one particular subject *i* is calculated the following:

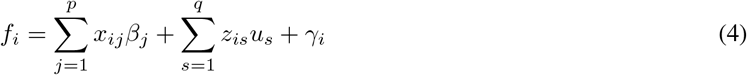

With

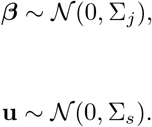

Here, ***β*** is a 1 × *p* vector containing the fixed regression weights corresponding to an *n* × *p* input matrix **X** with columns *j* = 1,. .., *p*. In case of non-centralized data one column of ones for an intercept offset has to be added. Similarly, **u** is a 1 × *q* vector containing the weights for random effects across subjects, corresponding to a dummy coded *n* × *q* matrix **Z** modeling site. For all linear models in Eq. 3 we assume *γ*_*i*_ = 0.

For the non-linear models, we assume *γ* is a Gaussian Process with mean function *m*(*x*) and covariance function *k*(*x,x*′) to allow for non-linear dependencies between the predictors and the target variable:

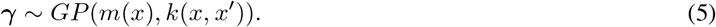

In our case, we set *m*(*x*) = 0 and define *k*(*x,x*′) as the additional non-linear component in the following squared-exponential form:

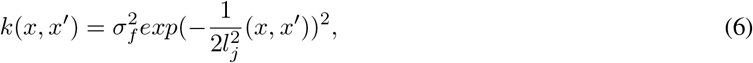

with free parameters for the signal variance term 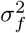 and the length scale *l*. Note this allows to specify two sources of variance: The signal variance 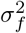 and the noise variance *σ*^2^ as modeled in Eq. 1.

From a hierarchical Bayesian point of view, random effects are equal to a hierarchical structure of sources of variation. For modeling site effects, introducing a hierarchical structure has the benefit that it allows to include structural dependencies between sites via partial pooling. Thus, instead of modeling site effects as an effect shared between sites or independently from each other, a semi-independent association between sites can be obtained via assuming that all site parameters originate from a shared first-order prior distribution. This concept has been used elsewhere (Kia et al., 2020; Gelman et al., 2013; Bonilla et al., 2008).

We hence induce shared priors and hyper priors ***θ***_**0**_ for site *s*, i.e. ∀_*S*_, *u*_*s*_ ∼ *Inv*Γ (2, 2), and a uniform prior for the length scale *l* ∼ *U*(1, 8). We use Stan (Carpenter et al., 2017; Stan Development Team, 2020b) to estimate all free parameters ***θ***= (***β***^***T***^, **u**^***T***^, ***l***^***T***^, *σ, σ*_*f*_) performing Bayesian inference:

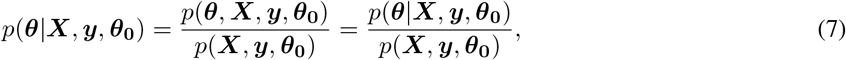

where *p*(***X, y, θ***_**0**_) = ∫*p*(***θ***)*p*(***X, y, θ***_**0**_|***θ***) *d****θ***.

#### 3.8.1 Posterior predictive distribution

We obtain the posterior predictive distribution *y*_∗_ for a new sample *x*_∗_ via:

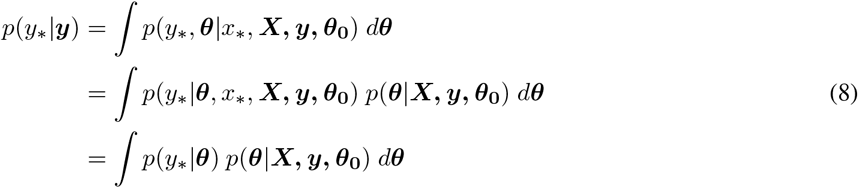

as ***y*** and *y*_∗_ are considered to be conditionally independent given ***θ*** (Gelman et al., 2013).

Further, the predictive distribution can be computed exactly, writing the joint distribution of the known data ***y, X*** and the new sample *x*_∗_, with the variance being determined by sample variance σ^2^ and the Gaussian kernel *k*(*x, x*′):

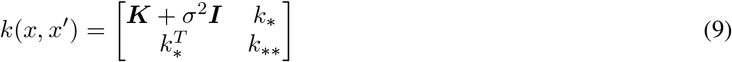

Here, ***K*** is an *n* × *n* covariance matrix of training data, *k*_* *_ denotes the variance at the test sample points and *k*_*_ is the covariance between *y*_*_ the known data.

Finally, each individual’s *z*-score of deviation can be calculated via:

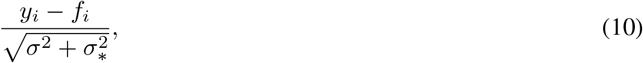

where 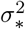 is the predictive variance that can be calculated the following (see also (Rasmussen and Williams, 2006)):

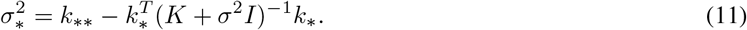

#### 3.8.2 Comparison models

We compare the hierarchical Bayesian attempt to normative modeling to commonly used harmonization techniques in which site is controlled for by subtracting an estimate of the site effect from the data prior to fitting the normative model. These methods included: i) removing additive effects of site, by regressing out site effects via linear regression and using the residuals as input for the simple Bayesian linear model to obtain the normative scores, ii) harmonizing for additive and multiplicative effects of site using ComBat (Johnson et al., 2007; Fortin et al., 2017), iii) *modified ComBat*, thus, using ComBat as before, but preserving biological variance of interest i.e., sex and age. All these methods involve removing site effects prior to estimating the normative scores in contrast to our method in which we explicitly model site within the normative modeling framework. These harmonized data, obtained as output from the harmonization techniques, are subsequently used for normative modeling in a simple Bayesian linear model that does neither take into account site effects nor non-linear dependencies between age and measures of cortical thickness. Thus, Eq. 3 is reduced to ***f* = *Xβ*** with ***β* ∼ 𝒩** (0, ∑_*j*_). In addition we use this simple Bayesian linear model to make one set of predictions for each regions from data that was not in any way harmonized for site (raw data model).

R (R Core Team, 2020) was used for preprocessing of all data and to create the data set where site was regressed out, and for preprocessing the data with ComBat (Johnson et al., 2007; Fortin et al., 2017).

#### 3.8.3 Implementation: Normative modeling in Stan

Both the hierarchical Bayesian and the comparison model version of the normative models were implemented in Stan (Carpenter et al., 2017; Stan Development Team, 2020b), a probabilistic C++ based programming language to perform Bayesian Inference, and analyzed in R (R Core Team, 2020) using the package rstan (Stan Development Team, 2020a). Stan allows to directly compute the log posterior density of a model given the known variables *x* and *y*. It uses the No-U-Turn Sampler (NUTS) (Hoffman and Gelman, 2014), a variation of Hamiltonian Monte-Carlo Sampling (Duane et al., 1987; Neal et al., 2011; Neal, 1994) to generate representative samples from the posterior distribution of parameters and hyper parameters *θ*, each of which has the marginal distribution *p*(*θ*|*y, x*). This is achieved by first approximating the distribution of the data to a defined threshold in a warm up period and then randomly sampling from the model, generating new draws of parameters for each iteration and calculating the response of the model. This approach of sampling instead of fitting allows for the simulation of complex models for which the derivation of an analytical solution of the posterior is computationally costly or not possible.

The Bayesian framework provides access to the full posterior distribution and to the distribution of all parameters. This allows to deduce the a variance estimate of each parameter, leading to a parameter estimate that is not only described by its mean, but also by the (un)-certainty around the mean estimation, providing information on its accuracy and reliability. Moreover, we can use the posterior distribution of each site-specific parameter from the training set as prior for the test set, allowing to make predictions for unfamiliar sites.

The Stan code for the HBLM, the HBGPM and the simple Bayesian linear model without site as predictor can be found at https://github.com/likeajumprope/Bayesian_normative_models.

### 3.9 Model simulation process in Stan

Parameters indicating the goodness of the model simulation process in Stan (Carpenter et al., 2017; Stan Development Team, 2020b) itself, like convergence, effective sample size and trace plots can be found in the supplementary material.

## 4 Results

### 4.1 Comparing hierarchical Bayesian models and comparison models

Both the HBLM and the HBGPM outperformed all other comparison models with respect to all performance measures considered in this study. In detail, the HBLM and the HBGPM showed higher average values of the Pearson’s correlation coefficient *ρ* (Table 2), lower average SRMSEs (Table 3), smaller average LL (Table 4) and higher average proportions of EV (Table 5) than all comparison models (p < 0.001 for all comparisons). For none of these comparisons did the non-linear HBGPM outperform the linear HBLM. In addition to the mean comparisons reported in Table 2 - 5, the distribution of all performance measures across all 34 regions and for average cortical thickness across the entire cortex per model can be found in Fig. 4. A detailed comparison of all models with respect to to *ρ*, SRMSE, EV and LL can be found in the supplementary material.

**Table 2:**
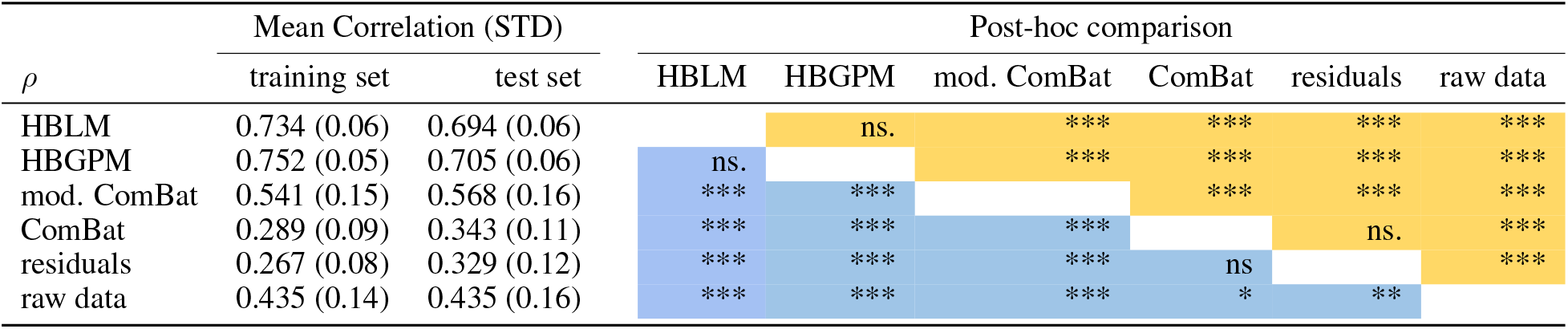
Post-hoc tests of correlations between true and predicted values. Cell values indicate post-hoc comparison significance values (adjusted by tukey method for a comparing a family of 6 estimates). Signif. codes: 0 ‘***’ 0.001 ‘**’ 0.01 ‘*’ 0.05 ‘.’ 0.1 ‘ ‘ 1 ns. blue: test set. yellow: training set.

**Table 3:** Post-hoc tests of SRMSE between true and predicted values. Cell values indicate post-hoc comparison significance values (adjusted by tukey method for a comparing a family of 6 estimates). Signif. codes: 0 ‘***’ 0.001 ‘**’ 0.01 ‘*’ 0.05 ‘.’ 0.1 ‘ ‘ 1 ns. blue: test set. yellow: training set.

| SRMSE | Mean SRMSE (STD) |  | Post-hoc comparison |  |  |  |  |  |
| --- | --- | --- | --- | --- | --- | --- | --- | --- |
|  | training set | test set | HBLM | HBGPM | mod. ComBat | ComBat | residuals | raw data |
| HBLM | 0.0608 (0.006) | 0.066 (0.005) |  | n.s | *** | *** | *** | *** |
| HBGPM | 0.0587 (0.006) | 0.064 (0.006) | ns. |  | *** | *** | *** | *** |
| mod. ComBat | 0.0763 (0.007) | 0.075 (0.008) | *** | *** |  | *** | n.s | ns. |
| ComBat | 0.0872 (0.003) | 0.085 (0.005) | *** | *** | *** |  | *** | *** |
| residuals | 0.0865 (0.003) | 0.085 (0.004) | *** | *** | ns. | *** |  | n.s |
| raw data | 0.0808 (0.006) | 0.085 (0.008) | *** | *** | *** | *** | *** |  |

**Table 4:** Averaged log loss for training and test set.

| LL | training set | test set |
| --- | --- | --- |
| HBLM | -1.050 | -1.121 |
| HBGPM | -1.020 | -1.109 |
| ComBat mod. | -1.225 | -1.193 |
| ComBat | -1.374 | -1.336 |
| residuals | -1.381 | -1.394 |
| raw | -1.299 | -1.335 |

**Table 5:** Averaged explained variance for training and test set.

| EV | training set | test set |
| --- | --- | --- |
| HBLM | 0.5674 | 0.5 |
| HBGPM | 0.5397 | 0.485 |
| ComBat mod. | 0.3146 | 0.338 |
| ComBat | 0.0918 | 0.122 |
| residuals | 0.0778 | 0.114 |
| raw | 0.2091 | 0.208 |

**Figure 4:**
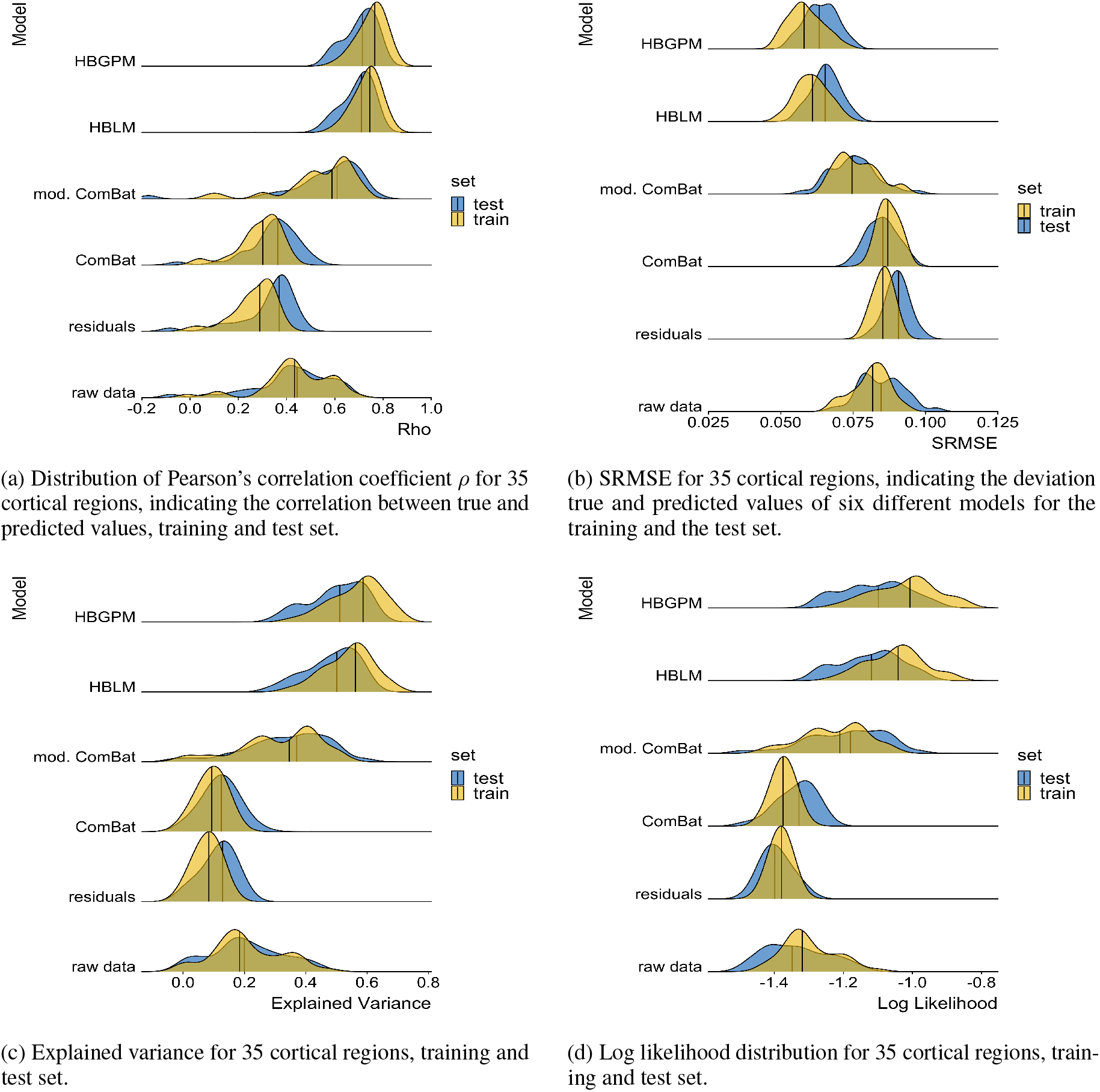
Performance measures.

#### 4.1.1 Mean standardized log-loss

To also account for the second order statistics of the posterior distributions created by each model, we calculated the mean standardized log-loss (MSLL). This measure can only be calculated for the test set, as it is the log loss standardized by the mean loss of the training data set (Rasmussen and Williams, 2006). Hence, the MSLL gives an indication of whether a model is able to predict the data better than the mean of the training set (with more negative values being better). An overview of the MSLL for all cortical thickness measures of all regions for all models is given in Fig. 5a. The only models that perform better for most regions than the mean of the training data set are the Hierarchical Bayesian models (*MSLL*_*HBGPM*_ < 0 for all regions; *MSLL*_*HBLM*_ < 0 for all but one region), in contrast to prediction from the *residuals* and the *ComBat* model, where none of the predictions perform better than the mean of the training data set (*MSLL*_*residuals*_ > 0 for all regions; *MSLL*_*ComBat*_ > 0 for all regions, see Fig. 5a. The MSLL for the *modified ComBat* model and *raw data* model were region-dependent, with 45 % regions (16 out of 35) for the *modified ComBat* model and 17% of regions (six out of 35) for the *raw data* model performing better than predictions from the mean of the training set. It should also be mentioned that for some individual regions the comparison models performed very poorly (max *MSLL*_*ComBat*_ = 356, max *MSLL*_*mod*.*ComBat*_ = 138, max *MSLL*_*raw*_ = 1252; max *MSLL*_*residuals*_ = 517) and show measures that exceeded the plotted range of Fig. 5a. In contrast, the maximum MSLL for the hierarchical Bayesian models was max -0.056 for the *HBGPM* and max 0.08 for the *HBLM*.

**Figure 5:**
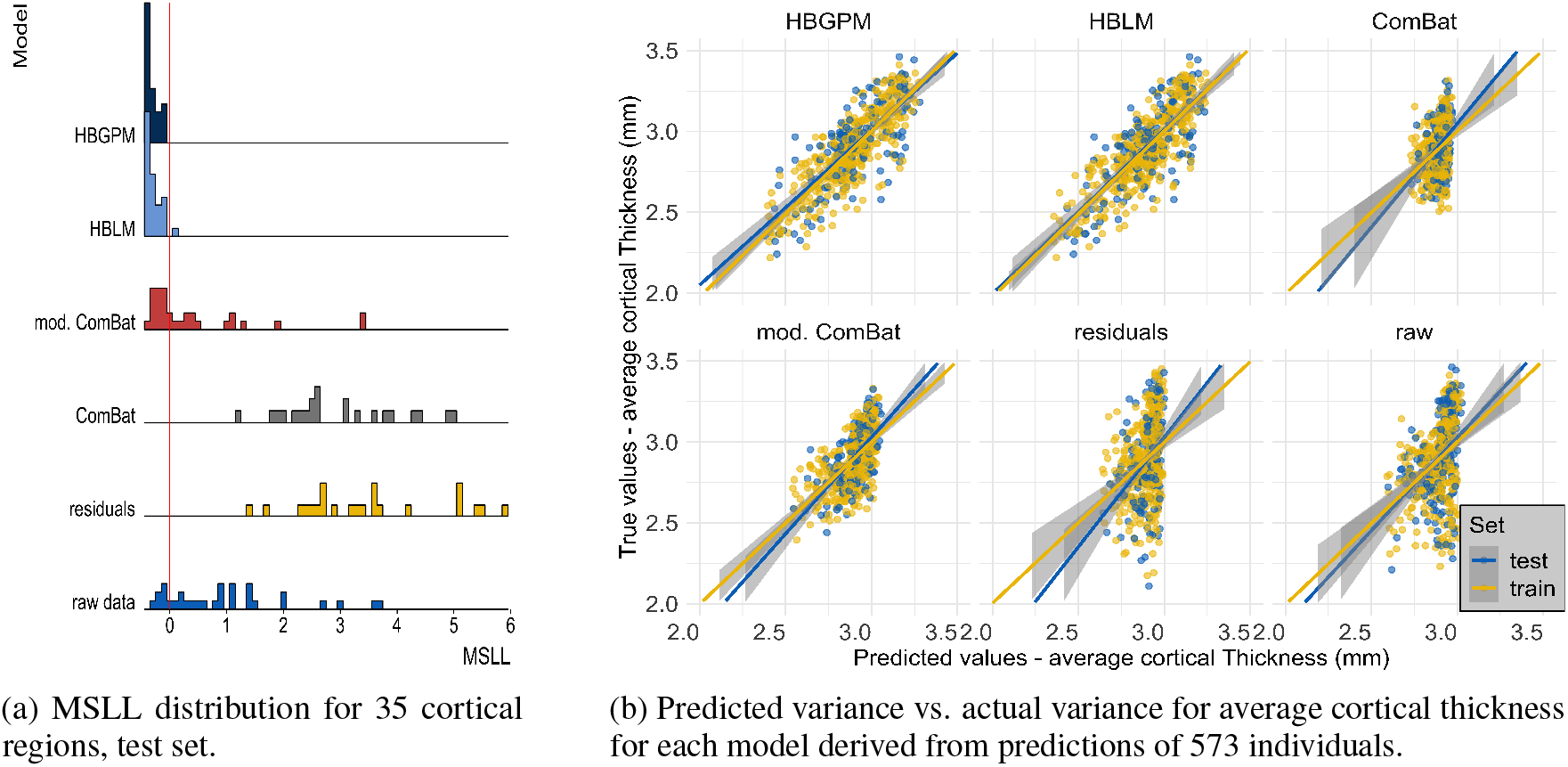
Mean standardized log loss and predicted variance for 35 cortical regions.

#### 4.1.2 Predictive Variance

We also observed that the models differ in the variance of predicted values, as visualized in Fig. 5b for average cortical thickness. For the *ComBat*, the *raw data* and the *residuals* model the range of predicted values was severely restricted (range predicted values *raw data*, test set: [2.60 - 3.03], range predicted values *residuals*, test set [2.64 - 3.00]; range predicted values *ComBat*, test set: [2.73 - 2.97]. These intervals cover 9.2 %, 7.9 % and 8.0 % of the original test set variance, respectively. The *modified ComBat* model retained 29.0% of the original test set variance (range predicted value *modified ComBat* [2.55 = 3.01]. In other words, all harmonization techniques had a reduced predictive variance and were instead biased toward predicting the mean. In contrast, this bias was substantially reduced in the hierarchical Bayesian models, which retained 57.0 % (*HBLM*) and 65.0 % (*HBGPM*) of the original test variance (range predicted values *HBLM*, test set: [2.43 - 3.23]; range predicted values *HBGPM*, test set: [2.38 - 3.28]). This highlights that the deviations from a two stage regression procedure where site effects are removed may be quite different from a model that accounts for age in a single regression model. In this case the residuals for both variants of Combat were considerably larger than *HBLM* and *HBGPM*.

### 4.2 Predicting site from z-scores

In order to test whether the HBLM and the HBGPM properly accounts for site, we subsequently analyzed whether site could still be predicted from the z-scores derived from those methods. An overview over the site effects before and after correction with HBLM and HBGPM is give in Fig. 6, in which the heterogeneity is visible reduced in corrected samples in panels 6d - 6f for the HBLM and 6g - 6i for the HBGPM (also note the difference in range of the x-axes between corrected and uncorrected plots). Using ANOVAs with site as predictor indicates that site cannot be predicted from the z-scores in the HBGPM control training set (p = 1) compared to p < 0.001 for the uncorrected training set, and is reduced in the HBGPM control test set (p = 0.03) compared to p < 0.001 in the uncorrected control test set, although site can still be predicted. For the HBLM, we find that the site cannot be predicted neither from the z-scores of the training set (p = 1) nor from the test set (p = 0.063). For the autism data set, we equally find that for the HBGPM the site effect is reduced for the z-scores both in the autism test set (p = 0.004) and compared to the uncorrected sets (p < 0.0001), as well as for the HBLM (test set: p = 0.01, uncorrected p = 0.0001), although still significant. In total, we observe that the linear model performs slightly better in correcting for site than the non-linear model.

**Figure 6:**
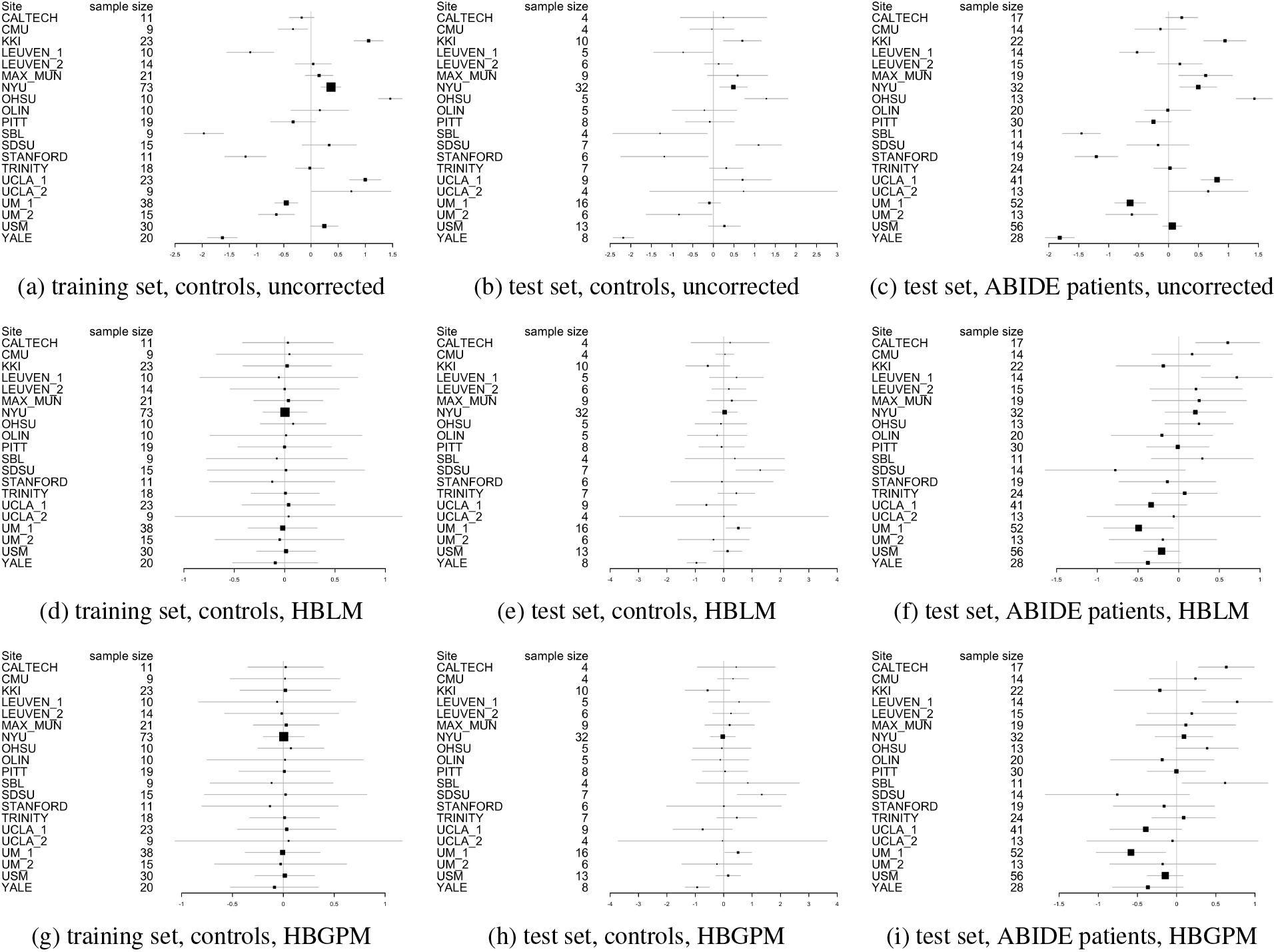
Forrest plots indicating the heterogeneity between sites. Uncorrected (6a - 6c), corrected with HBLM (6d-6f) and corrected with HBGPM (6g -6i). Also note the difference in range along the x-axes.

### 4.3 Application to clinical sample

It is desirable for any method that can accommodate unwanted site variance still preserves clinically relevant variance. In order to test whether our method is capable of retaining variance of interest, we applied the HBLM and HBGPM to the patient data set of the ABIDE sample and compared the clinical and the test control sample with respect to atypical z-values per region. We define a z-score of ± 1.96 based on the training set as atypical value, thus marking an individual that lies above or below the 95% percentile. As expected, for the control test set the average percentage of individuals with extreme z-scores is 5.7% for the HBGPM and 5.5% for the HBLM. For the autism sample, those numbers lie at 7.8% for the HBGPM and 7.2% for the HBLM, respectively. An overview over the distribution of percentages per region is given in Fig. 7 for controls and in Fig. 8 for the autism data set. The distribution of atypical z-scores across regions for both models is given in Fig. 9.

**Figure 7:**
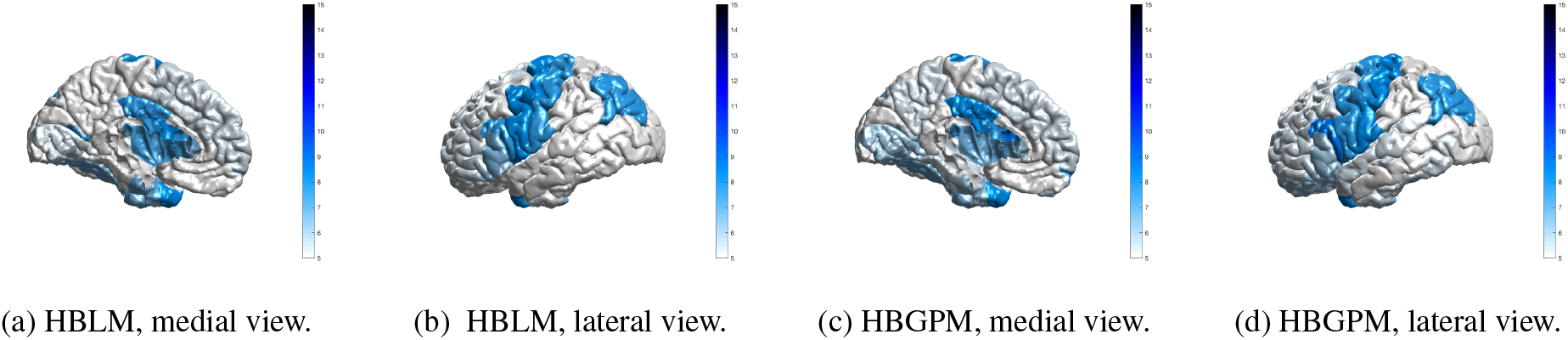
Region specific prevalence of atypical z-scores, control test set. Prevalence values of individuals with a z-score of ± 2SD, for the HBLM and the HBGPM model. Scores are thresholded at 5%, which is the expected amount of z-scores of ± 2SD within a normative model

**Figure 8:**
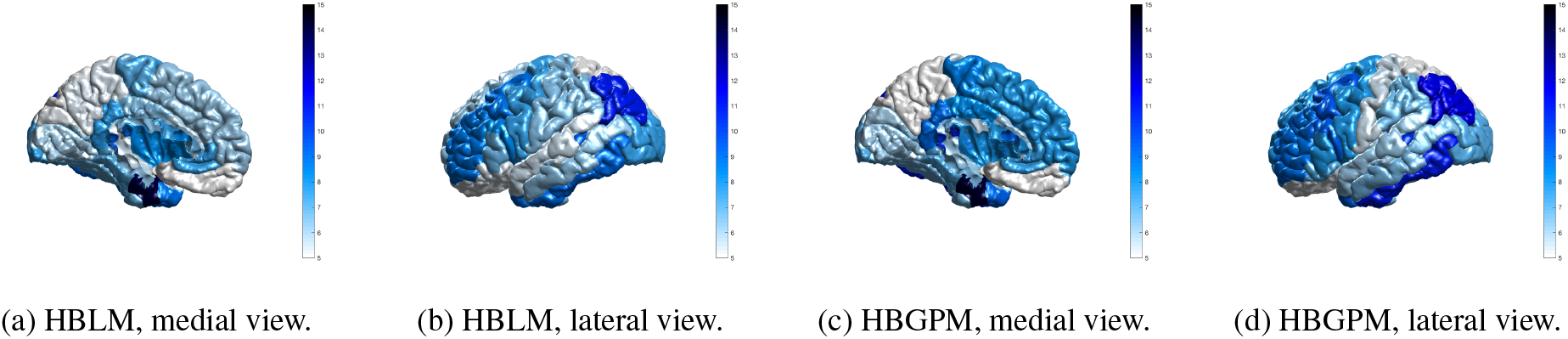
Region specific prevalence of atypical z-scores, autism test set. Prevalence values of individuals with a z-score of ± 2SD, for the HBLM and the HBGPM model. Scores are thresholded at 5%, which is the expected amount of z-scores of ± 2SD within a normative model

**Figure 9:**
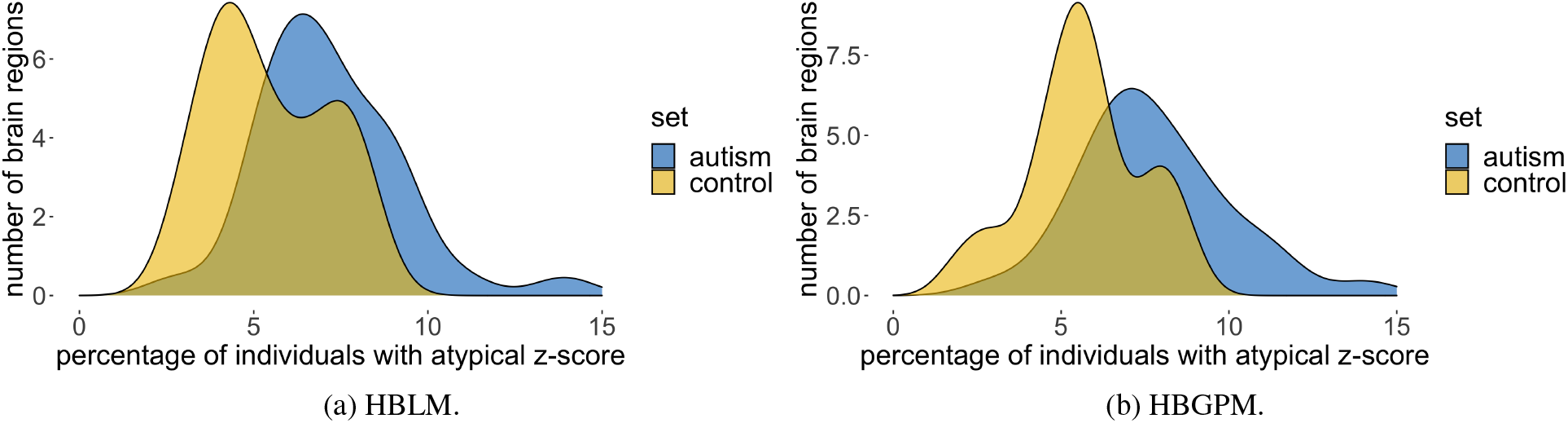
Prevalence of atypical z-scores across all regions.

## 5 Discussion

In this work, we aimed to provide a method that allows the application of normative modeling to neuroimaging data sets that are affected by site effects resulting from pooling data between sites. In contrast to other methods of harmonizing for additive and multiplicative site effects in the data prior to the normative modeling (e.g., regressing out site effects, harmonization with *ComBat*), our approach is based on modeling site as predictor within the normative modeling framework in a single regression model. The benefit of this approach is that it does not entail removing variance and thus cannot lead to an accidental removal of meaningful variation in case variables of intereest are correlated with site variation. Using a hierarchical Bayesian approach, we proposed two versions of normative models that were able to accommodate site effects. In both versions, site was modeled via a random intercept offset, but one version only models linear effects of age on cortical thickness (Hierarchical Bayesian Linear Model, HBLM), whereas the other version also included a Gaussian process term in order to allow potential non-linear relationships between age and cortical thickness measures (Hierarchical Bayesian Gaussian Process Model; HBGPM).

The normative models were trained on a training set consisting of healthy individuals from the ABIDE data set (70% of the data from 20 different sites, within-site split, preserving the distribution of age and sex across training and test set) and we presented results from generalization to a test set (the remaining 30% of the data from the same sites). We compared the performance of our hierarchical Bayesian normative models explicitly modeling site effects applied to cortical thickness measures derived from FreeSurfer (Fischl et al., 2004)) to other commonly used methods to deal with site effects. These alternative methods included: i) regressing out site via linear regression and using the residuals, removing additive site variation, ii) applying *ComBat* (Fortin et al., 2017, 2018) to harmonize additive and multiplicative site effects in the data, and iii) *modified ComBat*, hence applying ComBat while preserving age and sex effects in the data. Cortical thickness measures cleared from site effects using these alternative methods were used as dependent variables in a normative model with age and sex as predictors but excluding site. For comparison reasons, we also included a fourth model where we made predictions from raw data uncorrected for any site effects. Subsequently, we applied the hierarchical Bayesian models trained on healthy controls to the autism sample of the ABIDE data set.

We reported three main findings: (1) Our normative hierarchical Bayesian models (both the linear *HBLM* and non-linear *HBGPM* versions), explicitly modeling site effects within the normative modeling framework, outperformed all alternative harmonization models with respect to model fit, including correlations between true and predicted values (*ρ*), standardized root mean square error (SRMSE), explained variance scores (EV), log-likelihood (LL) and the mean standardized log-loss (MSLL); (2) the non-linear model did not significantly improve prediction of cortical thickness based on age, sex and site compared to the linear model; (3) all methods, but in particular the harmonization comparison methods lead to an undesirable shrinking of the variance in the predictions. We subsequently showed that in the z-scores obtained from the hierarchical Bayesian model site variation is substantially reduced, while retaining clinically useful variation.

We showed that when using neuroimaging structural data sets pooled across different sites and scanners for estimating normative models, better predictive performance could be achieved by including site as a predictor than using a two-step approach of first harmonizing the data with respect to site and subsequently creating a normative model using these “cleared” data. This conclusion was based on results showing that the hierarchical Bayesian models outperformed the harmonizing comparison models on all of the performance metrics we examined. This included the predictions derived from data that was cleared from site effects by a version of ComBat (Fortin et al., 2017, 2018) in which variation associated with age and sex was preserved, which was the best performing method across all harmonizing comparison models. We observed a higher correlation between true and predicted values and LL values closer to zero for our hierarchical Bayesian models explicitly modeling site effects with a random intercept offset, indicating better model fit. As a key factor of normative models is that they are not only able to estimate the predictive mean, but also give an estimate of the predictive variance and variation around the mean (Marquand et al., 2019, 2016), we also included explained variance scores and the MSLL as performance metrics. Our HBLM and HBGPM models showed higher explained variance than the alternative models. In addition, the HBLM and HBGPM showed a negative MSLL in the test set; a metric which contrasts the log-loss between the true and predicted values by the loss that would be achieved using the mean and the variance of the training set (Rasmussen and Williams, 2006), thus capturing differences in variance in the data sets. This benefit in performance for the hierarchical Bayesian models is in line with previous literature using a similar paradigm (Kia et al., 2020). Kia et al. (2020) showed that a hierarchical Bayesian regression approach using site as a batch effect lead to a better performance than complete pooling, no-pooling and ComBat. In detail, our findings match Kia et al. (2020)’s findings with respect to the comparison between a normative model created from hierarchical Bayesian regression (HBR) and a *modified ComBat* version in a data set with the same sites in training and test set. Their findings are in line with ours with respect to *ρ* ((Kia et al., 2020): *HBR* range: 0.4 - 0.9, *modified ComBat* range: 0.2 - 0.8), SMSE: ((Kia et al., 2020): *HBR* range: 0.2 - 0.9, *modified ComBat* range: 0.4 - 1.0) and MSLL ((Kia et al., 2020): *HBR* range: -0.7 - -1.0, *modified ComBat* range: -0.04 - 0.0), except that the MSLL for the *modified Combat* model was worse in our study (see Figs. 4a, 4b, 5a). Therefore, our findings replicate the findings of Kia et al. (2020) using an independent data set and separate implementation and extend that method to model non-linear functions using a Gaussian process term.

We anticipated that the non-linear version of the normative model, which included a Gaussian Process for age, would perform better than the linear version, as studies have shown that the association between age and regions of cortical thickness can be non-linear, especially for older age ranges (Storsve et al., 2014). However, our results showed similar performance in predicting cortical thickness based on age, sex and site for both linear and non-linear models. This might be due to the fact that the the age range in our sample was restricted, ranging from 6-40 years, thus likely capturing an age range where the association between age and cortical thickness is still mostly linear (Wierenga et al., 2014). As a consequence, the non-linear version of the model was not able to improve the overall performance. Nonetheless, since other structural brain measures, including sub-cortical volumes and cortical surface area (Wierenga et al., 2014; Raznahan et al., 2011), have shown stronger non-linear associations with age, non-linear normative models may outperform a linear model for other types of structural brain imaging measures.

Despite an overall good performance of our models, it should also be mentioned that the performance showed substantial variation between regions, as reflected in the variation in *ρ* values, SRMSE, EV, LL and MSLL within models. We assume that this due to the fact that, although *average* cortical thickness shows a strong association with age, different cortical brain regions differ in their association with age and the magnitude of this correlation also changes across the lifespan ((Storsve et al., 2014)).

All models, but in particular the comparison models, had a significant shrinkage effect on the variance of the predicted values, indicating that harmonization techniques remove variance that is useful in predicting the response variable. This was most extreme for regressing out site effects and lead to poor performance across all performance metrics. We also observed that the performance of the *residuals* model was similar to the *ComBat* model without the preservation of age and sex, which was particularly reflected in the similarities of predicted variance in Fig. 5b and in the SRMSE. More importantly, both models suffered a loss of more than 90% of their original test variance. This is evident in 4 in that the range of the predicted values is less than the range of the true values. This is particularly relevant for normative modelling where the primary interest is in quantifying deviations from a reference model via individual level Z-statistics. Since the model residuals are different for models that regress out site effects in advance, it is clear that these will yield different deviations and different downstream inferences. In the case shown in (Fig. 4), the model residuals have a higher magnitude because the model explains less variance in age. In contrast, the performance improved when variables like age and sex were preserved, as demonstrated by an increase in performance measures when using the version of *ComBat* in which variation associated with age and sex was preserved. We argue that the similarity in performance between *ComBat* and the *residuals* model is an indicator of the same underlying process, showing a weakness of the harmonization approach: merely regressing out site effects led to the removal of meaningful variation correlated with the predictors of interest (in this case age and sex), especially when these predictors of interest were correlated with the site effects, which subsequently led to worse predictions of cortical thickness based on age and sex. This could be partially prevented by preserving important sources of variation when regressing out site effects, as shown for the *modified ComBat* model, where specified sources of variance were preserved when regressing out site effects. However, our results showed two additional flaws of the harmonization approach: 1) as already pointed out by Kia et al. (2020), in order to specify sources of variance that should be retained, all those sources of variance have to be known, which is not always the case; 2) even with age and sex preserved the modified ComBat model only retained 40% of the original variance. Our hierarchical Bayesian models including the prediction-based approach, in contrast, preserved known and unknown interactions between site and biological covariates by specifically modeling site, thus overcoming this requirement. The result was reflected in larger proportions of variance retained (see Fig. 5b. The advantage of the hierarchical Bayesian approach becomes particularly clear when considering that the scores derived from normative models are relative scores describing the deviation from a predicted normative mean. Thus, the normative deviation score is not affected by the absolute value of the predicted mean, and the number of predictors in the model does not influence the normative score.

Previous attempts to estimate the centiles of normative models have included polynomial regression (Kessler et al., 2016), support vector regression (Erus et al., 2015), quantile regression (Huizinga et al., 2018; Lv et al., 2020) and Gaussian process regression (Wolfers et al., 2018b), providing different degrees of the ability to separate between sources of variances and making individual predictions (for an overview see (Marquand et al., 2019)). We chose a hierarchical Bayesian framework for the implementation of our normative model as it has several advantages. The distribution-based structure based on posteriors allows for the separation and integration of different sources of variances, including epistemic (uncertainty in the model parameters), aleatoric (inherent variability in data) and prior variation, which are all considered when predicting cortical thickness based on age, sex and site. This allows for both the integration of already known information in the form of priors into the predictions, and for an adjustment of the precision of the estimate based on the uncertainty at each data point. In addition, the Bayesian framework, as implemented in software packages like Stan (Carpenter et al., 2017; Stan Development Team, 2020b), allows to draw samples from the full posterior distribution at the level of individual participants, which leads to an exact estimate of all parameters instead of an approximation. In particular in comparison to quantile regression, the distributional assumption entailed in the hierarchical Bayesian approach also allows to get more precise estimates of the underlying centiles, particularly in the outer centiles, which are usually of primary interest and where the data are sparsest. The proposed Bayesian framework also offers an elegant way to integrate site effects into normative models. Site effects can be modeled via a hierarchical random effect structure, in which different sites are modeled semi-independently, sharing variation via a combined prior of higher order. This approach, also known as partial pooling, allows for including site-specific variance into the prediction for site, while at the same time constraining the amount of between-site variation to a maximum.

Whilst the primary aim of this study was to develop a novel method for dealing with site effects specifically within a normative modeling framework, our results show that the method can be used as a general approach to clear neuroimaging data from site, age and sex effects, demonstrating no residual site effects in the training sets and substantially reduced residual site variance in the z-scores in the control and autism test sets. This can be explained by the fact that the normative score describes an individual’s cortical thickness *in relation to* the variance explained by the predictor variables in the normative model (age, sex and site). Hence, they can be seen as “cleaned” cortical thickness measures that can be the basis for further analysis, for example to establish the association between cortical thickness measures and clinical or demographic information. We also observed that the HBLM performs slightly better than the non-linear version in controlling for site effects, a finding that might be due to its ability to pick up noise variation between training and test which might particularly affect smaller sites. Further, we showed a difference in the distribution of atypical z-scores between the test control and the test autism set. The average percentage of atypical z-scores in our study for the autism sample was 7.8% for the HBGPM and 7.2% for the HBLM, compared to 5.7% for the HBGPM and 5.5% for the HBLM for the control test set. These findings illustrate, on the one hand side, that the models were both able to produce the per-definition expected amount of 5% of atypical z-scores in a healthy control test set, thus validating the model. On the other hand side, and more importantly, the difference in atypical z-scores between test healthy controls and the autism sample showed that and that the models are yet able to preserve clinically significant variation while removing site related variation. Both the regions affected by a large amount of atypical z-scores in autism and the percentage of individuals with atypical z-scores reported 9 are broadly in line with previous findings (Bethlehem et al., 2020; Zabihi et al., 2019b). However, due to the existence of laterality effects in autism (Jiao et al., 2010; Khundrakpam et al., 2017), which were likely concealed by averaging left and right hemisphere in our study, we refrain from discussing those results further. For the interested reader, discussions of the clinical implications of the ABIDE data set, yet without considering site effects can be found elsewhere (Bethlehem et al., 2020; Zabihi et al., 2019b).

Our proposed method has three potential disadvantages. The first one is related to the computational cost associated with estimating the covariance matrix within the Gaussian Process for the non-linear models, which in our analysis amounted to 25 hours per model per region and could only be mastered via parallel processing on a cluster. This is due to the fact that using the non-linear Gaussian Process term becomes very time and memory expensive with growing *n* (*O*(*n*^2^)). Thus, in cases in which the relationship between the predictor and the outcome is estimated to be close to linear, the need for the more complex non-linear model should be carefully considered. Secondly, the between-site split and the model at its current state only allow generalizations to a test set which includes individuals from the same sites as the training set, thus where the site variation is known. However, especially in clinical settings, generalizing the model and making predictions in data from new sites is an important additional goal. Despite the fact that we cannot use the posterior distribution of one particular site as a prior when applying the model to a new, unknown site, the hierarchical Bayesian framework still allows using the posterior parameter distributions of *all* sites as derived from the training data set as priors for site parameters when applying the model to a new site. This approach has already been successfully demonstrated in Kia et al. (2020) where the posterior parameter distribution of site derived from the training data was fed as a informative prior for the site predictor in a normative model applied to the test data consisting of new (unknown) sites. This use of a so called informed priors leads to more accurate and precise predictions than the broad, unspecific prior that would have to be used in cases where the distribution of the data is unknown (Kia et al., 2020). Thus, despite some loss in precision, the Bayesian framework can, in contrast to all other methods examined in this paper, be adapted to make predictions to new, unknown sites. Thirdly, this approach does not account for correlations between different brain measures, although we consider that a potential future extension

## 6 Conclusion

We proposed an extended version of a normative modeling approach that is able to accommodate for site effects in neuroimaging data. The method is superior to previous approaches, including regressing out site and versions of ComBat (Fortin et al., 2017; Johnson et al., 2007) and facilitates the estimation of normative models based on neuroimaging data pooled across many different scan sites, while retaining useful clinical variation. A further extension of the model to make generalizations to new sites and the application to clinical data will be the objectives of future work.

## 8 Data Availability Statement

We declare that all software, data and code used for this paper is publicly available. The ABIDE (autism brain imaging data exchange) (Di Martino et al., 2014) data setis available at: http://preprocessed-connectomes-project.org/abide/. The software package Stan (Stan Development Team, 2020b) is available at: https://mc-stan.org/users/interfaces/. The software package R (R Core Team, 2020) is available at: https://www.r-project. org. The R-package rstan (Stan Development Team, 2020a) is available at: https://cran.r-project.org/web/ packages/rstan/index.html. The Stan code for the HBLM, HBGPM and simple Bayesian linear model are available at: https://github.com/likeajumprope/Bayesian_normative_models.

## 9 Acknowledgment

LS was supported by the NHMRC Career Development Fellowship (1140764) and NIH RO1 (MH117601). AM grateful acknowledges funding from the Dutch Organisation for Scientific Research (NWO) under a Vernieuwingsimpuls VIDI fellowship (grant number 016.156.415), the European Research Council (consolidator grant, number 101001118) and the Wellcome Trust under a Digital Innovator grant (215698/Z/19/Z)

