## supplementary material for "Accommodating site variation in neuroimaging data using normative and hierarchical Bayesian models"

### 1 Supplementary material

#### 1.1 Model convergence, effective sample size and $\hat{R}$

Stan allows for the computation of a number of diagnostics on the quality of the Markov Chain Monte Carlo (MCMC) sampling process.

##### 1.1.1 Model convergence

Markov chains are defined to only generate samples from the target distribution after the distribution has converged to an equilibrium, thus, when the distribution is considered to be the target density. In theory, this equilibrium can only asymptotically be reached, as the number of draws is theoretically infinite. In practice, the number of draws has to be a-priori set to a finite amount. As a consequence, the actual and convergence to the target density has to be monitored [1, 2, 5]. One way to monitor the convergence of a chain to equilibrium is to compare the chain to other randomly initialized chains. This can either be done via visual inspection, or using the scale reduction statistic, $\hat{R}$ [3]. For visual inspection, the trace plots over 4 chains for all parameters can be found in Figs. 1a, 2a , 3a, 4a, 5a, 6a. The $\hat{R}$ values, indicating the convergence of chains, can be found in Figs. 1b, 2b , 3b, 4b, 5b, 6b for each model, respectively. All $\hat{R}$ values are <1.05, which provides good evidence that all chains have reached convergence and can therefore be considered to provide unbiased samples from the target density.

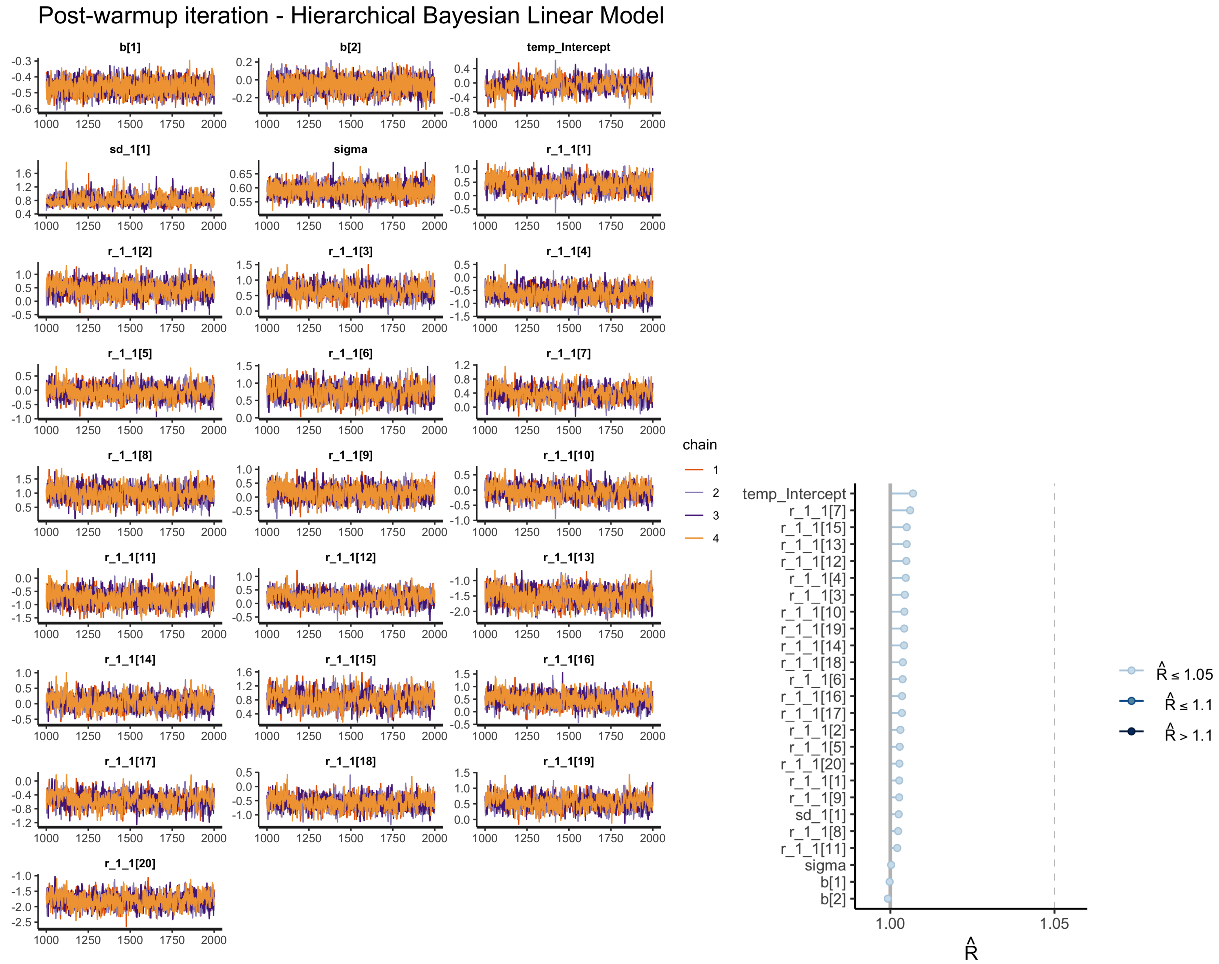


(a) Traceplots

(b) $\hat{R}$

Figure 1: Hierarchical Bayesian Linear Model

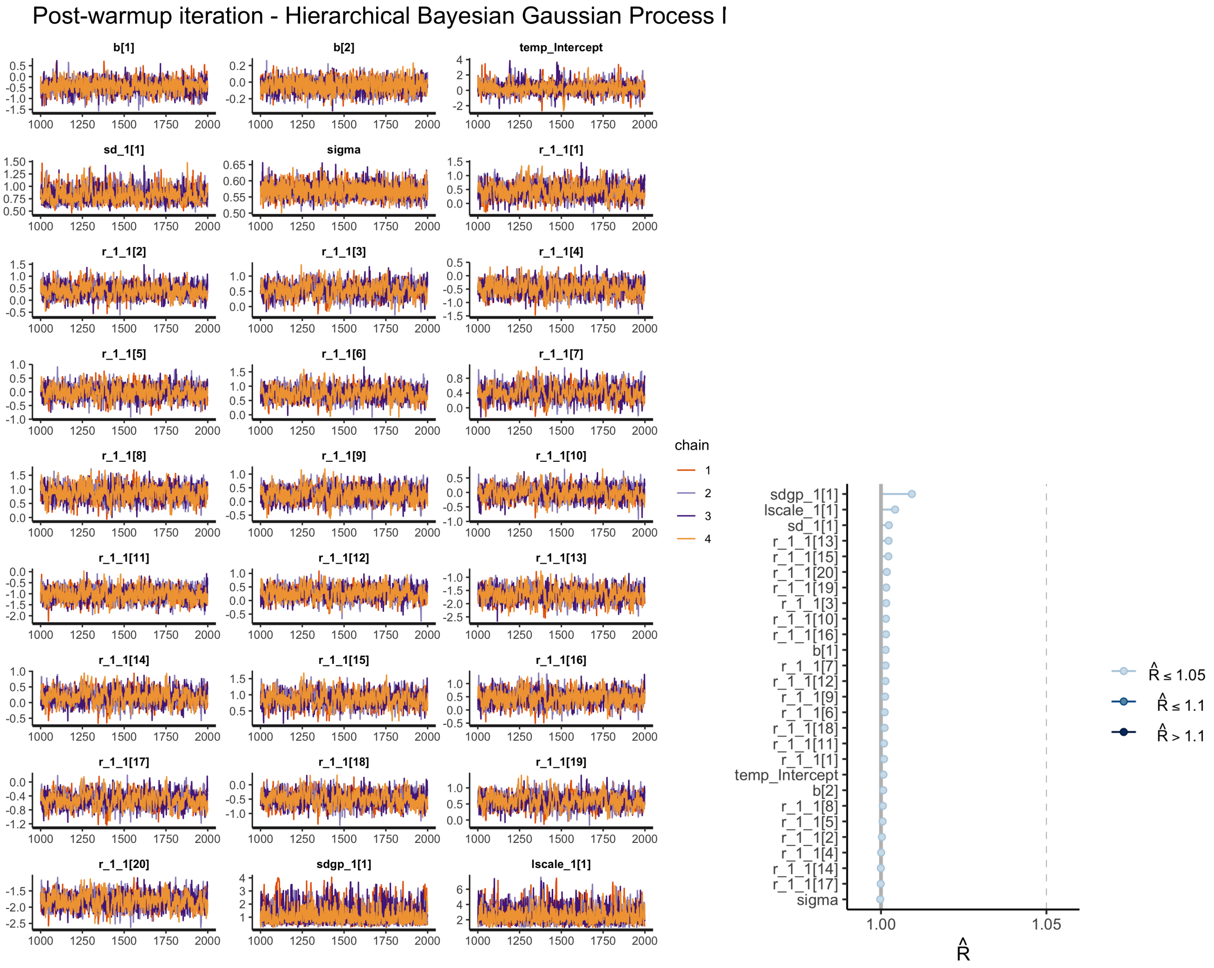


(a) Traceplots (b) $\hat{R}$

Figure 2: Hierarchical Bayesian Gaussian Process Model

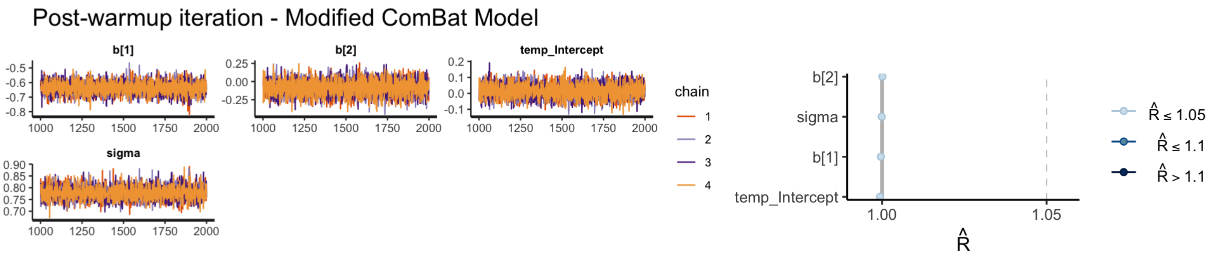


(a) Traceplots (b) $\hat{R}$

Figure 3: modified ComBat Model

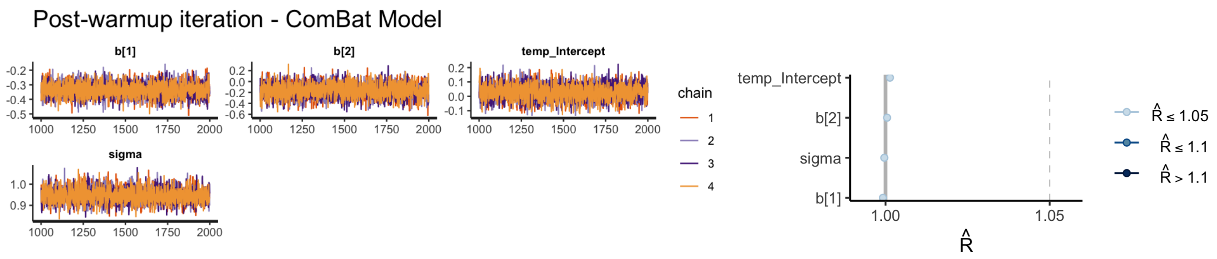


(a) Traceplots (b) $\hat{R}$

Figure 4: ComBat Model

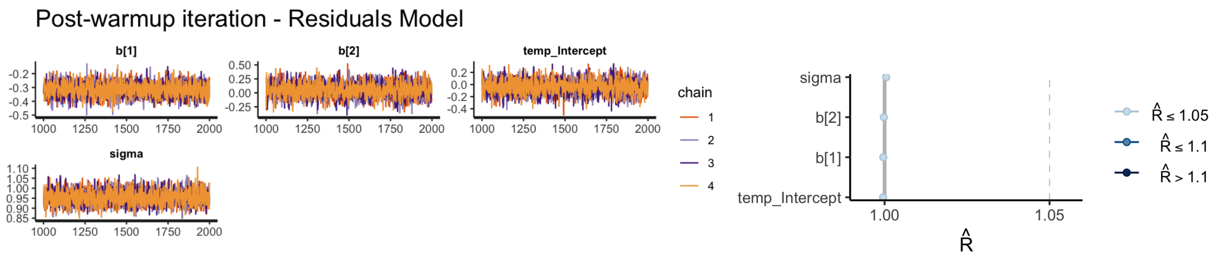


(a) Traceplots (b) $\hat{R}$

Figure 5: Residuals Model

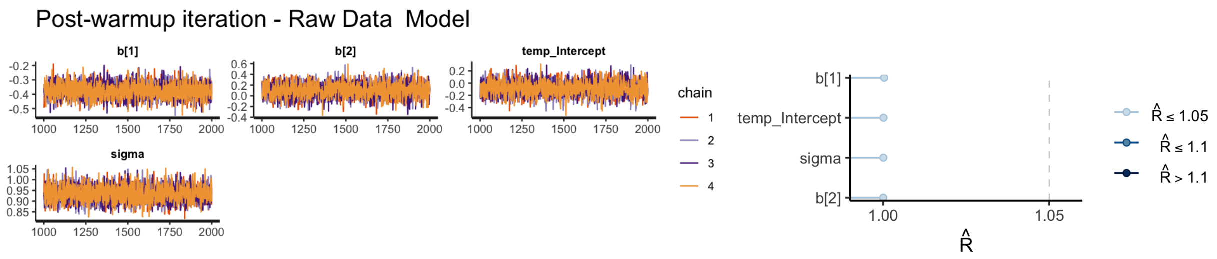


(a) Traceplots (b) $\hat{R}$

Figure 6: Raw Data Model

###### 1.1.2 Effective sample size

One characteristic of MCMC methods is that samples will be auto- or anti-correlated within a chain, leading to a reduction of precision in the estimates of posterior quantities [4]. Stan uses the auto correlation $\rho_t$ between samples $n$ and $n$+$t$ with lag $t$ to estimate the effective samples size $N_{eff}$ of independent samples in the chain. $N_{eff}$ is considered to have the same estimation power as $N$ correlated samples and is then used, rather than $N$, to estimate precision and error measures [5]. $N_{eff}$ for all models can be found in Figs. 7a, 7b, 7c, 7d, 7e, 7f.

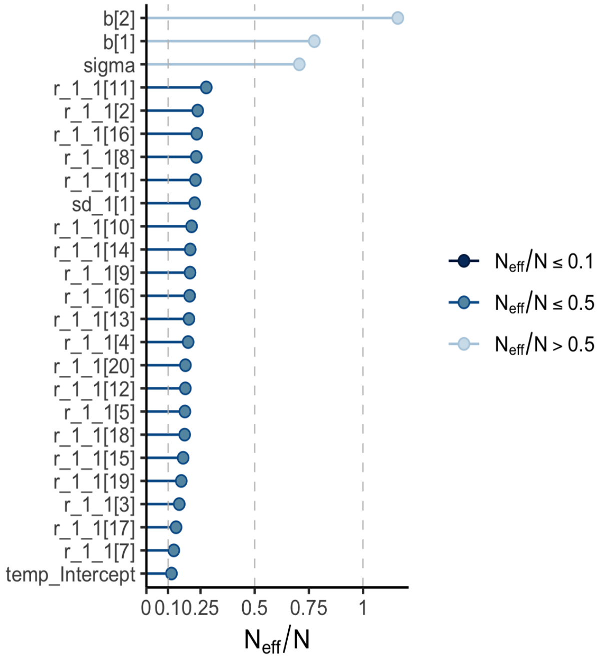


(a) Hierarchical Bayesian Linear Model

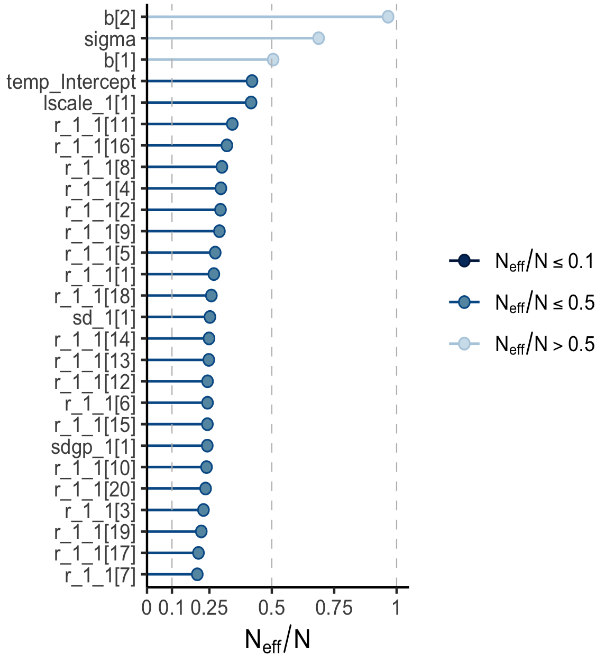


(b) Hierarchical Bayesian Gaussian Process Model

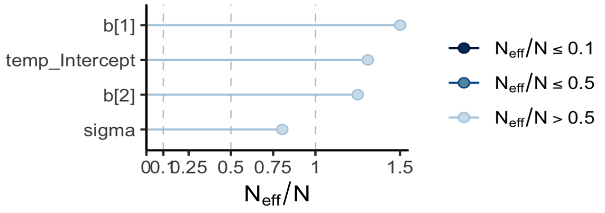


(c) Model ComBat.

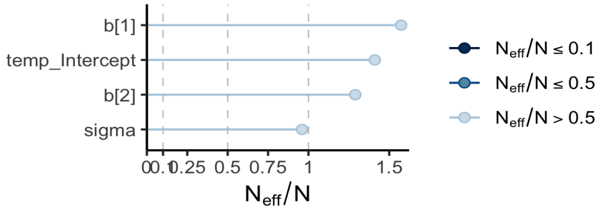


(d) Model modified ComBat w *age*/*sex* preserved

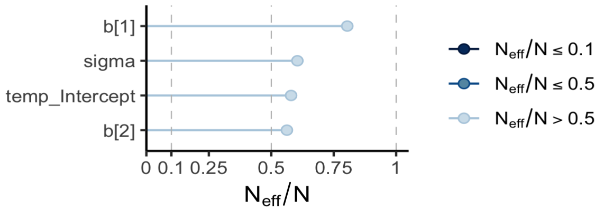


(e) Residuals Model

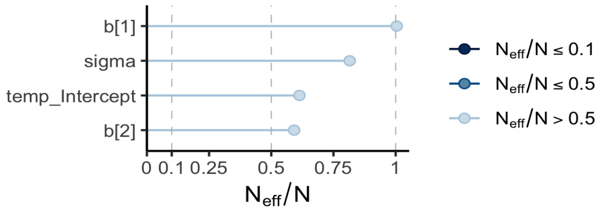


(f) Raw data Model

Figure 7: Effective sample sizes $N_{eff}$ for all parameters

##### 1.2 Results

##### 1.3 Correlation between true and predicted value

Correlations between true and predicted values for the HBLM and HBGPM are expressed in terms of the correlation coefficient $\rho$, calculated separately for each region. $\rho$ ranged from 0.60 to 0.84 in the training and 0.55 to 0.80 in the test set. Overall, for just the Bayesian models, correlations were higher in the training set and dropped in the test set (training set, across all regions: $\bar{\rho}_{HBLM}$ = 0.73, *SE* = 0.05; $\bar{\rho}_{HBGPM}$ = 0.75, *SE* = 0.06; test set: $\bar{\rho}_{HBLM}$ = 0.69, *SE* = 0.06; $\bar{\rho}_{HBGPM}$ = 0.69, *SE* = 0.06) [F(1,136) = 18.817, p < 0.0001)]. Correlations did not differ significantly between the HBLM and HBGPM. [F(1,136) = 2.158, p = 0.144].

Comparisons to the other, non-Bayesian models showed that true values of cortical thickness could be predicted significantly better with our models that included *site* as a predictor than with all other models in which site was regressed out prior to running the normative models, both for training and test set (t-test *HBLM* and any other model p< 0.001, t-test *HBGPM* and any other model p<0.001, both for training and test set.) The full distribution of the correlation coefficient $\rho$ for all 35 regions per model can be found in Fig 3a, main text. In addition, a test comparing the performance of all models for all regions (Bayesian and non-Bayesian) showed that predictions made from training data were not more accurate than predictions from the test data (interaction *set* $\times$ *model*, F[1,478] = 2.72, p = 0.020). Further inspection showed that this might have been caused by the *residuals*, the *ComBat* and the *modified ComBat* model, where the test set performed *better* than the training set, canceling out performance benefits of the training data in the HBLM and HBGPM (see also Fig. 3a, main text).

##### 1.4 Standardized Root Mean Squared Errors

We further evaluated the fit of the models by calculating the Standardized Root Mean Squared Error (SRMSE) between true values and predicted values per model per region. As expected, the SRMSE was larger for the test set (*M* = 0.078) and smaller for the training set (*M* = 0.075, [F(1,116316)=37.54, p < 0]). For both the training and the test set, the Bayesian models showed smaller SRMSEs than all other models across all regions (p < 0.001; training set: $SR\bar{M}SE_{HBGPM}$ = 0.059, *SE* = 0.0007; $SR\bar{M}SE_{HBLM}$ = 0.061, *SE* = 0.0007; $SR\bar{M}SE_{residuals}$ =0.087, *SE*=0.0007; $SR\bar{M}SE_{modifiedComBat}$ = 0.076, *SE* = 0.0007, $SR\bar{M}SE_{ComBat}$ = 0.087, *SE* = 0.0007; $SR\bar{M}SE_{rawdata}$ = 0.081, *SE*=0.007; test set: $SR\bar{M}SE_{HBGPM}$ = 0.064, *SE* = 0.001; $SR\bar{M}SE_{HBLM}$ = 0.066, *SE* = 0.001; $SR\bar{M}SE_{residuals}$ =0.087, *SE* = 0.001; $SR\bar{M}SE_{modifiedComBat}$ = 0.075, *SE* = 0.001; $SR\bar{M}SE_{ComBat}$ = 0.085, *SE* = 0.001; $SR\bar{M}SE_{rawdata}$ = 0.085, *SE*=0.001). Neither in the training nor the test set did the Bayesian models differ from each other (training set: contrast *HBLMR* - *HBLM*, t = 2.325, p = *ns.*; test set: contrast *HBLMR* - *HBLM*, t = 1.135, p = *ns.*). We also observed that both in the training and test set, the SRMSE of *ComBat* did not differ from the SRMSE of the *residuals* (training set: contrast *combat* - *residuals*, t = 0.690, p = *ns.*; test set: contrast *combat* - *residuals*, t = -1.703, p = *ns.*. The full distribution of SRMSE for all 35 regions per model can be found in Fig 3b, main text.

##### 1.5 Explained variance

Analysis of the proportion of variance explained EV = $\frac{\sigma^2_{\hat{y}-y}}{\sigma^2_y}$ were in line with the results reflected in $\rho$ and SRMSE. EV was higher for the *HBLM* and *HBGPM*, with an average of 0.56 (*HBGPM*, range: 0.35-0.70) and 0.53 (*HBLM*, range 0.35 - 0.67) for the training set and 0.50 (*HBGPM*, range 0.31 - 0.63) and 0.48 (*HBLM*, range 0.28 - 0.60) for the test set across all cortical regions. The proportion of explained variance was substantially lower for the comparison models, with the modified ComBat model performing best out of the comparison models, with an average of 0.31 for the training set (range: 0.00 - 0.51) and 0.33 for the test set (range -0.02 - 0.58) across cortical regions, but showing lower EV than the Bayesian models. Predictions derived from *residuals* and *ComBat* showed even lower EV, with *residuals* explaining an average of 0.07 for the training set (range: 0.00 - 0.15) and 0.11 for the test set (range 0.00 - 0.20) across cortical regions, and *modified ComBat* explaining an average of 0.09 for the training set (range: 0.00 - 0.17) and 0.12 for the test set (range: 0.00 - 0.25) across cortical regions. Thus, the *ComBat, modified ComBat and residuals* model performed even worse than predictions derived from *raw data*, which showed an average EV of 0.21 in the training set (range: 0.00 -0.46) and 0.20 in the test set (range 0.00 - 0.44) across cortical regions. These results include the interesting finding that the test set shows slightly higher EVs than the training set for all comparison models. An overview over the distribution of explained variance for training and the test set for all 35 regions for all models can be found in Fig. 3c, main text.

##### 1.6 Log likelihood

The point-wise log likelihoods (LL) between the true and predicted were calculated for each data point, summed up per model across regions and averaged y the number of individuals in training and test set, respectively. The averaged summed LL across regions was closer to zero for the nonlinear Bayesian model than for the linear Bayesian model, both for the training and the test set ( $\sum \frac{1}{n_{test}} LL_{HBGPM}$, test set: -1.109, $\sum \frac{1}{n_{test}} LL_{HBLM}$ test set: -1.121; $\sum \frac{1}{n_{train}} LL_{HBGPM}$, training set: -1.020, $\sum \frac{1}{n_{train}} LL_{HBLM}$, training set: -1.05. LL values were less close to zero for all comparison models, with the *modified Combat* model performing best for among those models, followed by the *raw data* model, the *residuals* model and the *ComBat* model (an overview of the log likelihood for all models is given in Tab. 4, main text) The distribution of the log likelihood for all regions is given in Fig. 3d, main text.
